# Insectivorous birds and bats outperform ants in the top-down regulation of arthropods across strata of a Japanese temperate forest

**DOI:** 10.1101/2023.11.07.565745

**Authors:** Elise Sivault, Jan Kollross, Leonardo Re Jorge, Sam Finnie, David Diez-Méndez, Sara Fernandez Garzon, Heveakore Maraia, Jan Lenc, Martin Libra, Masashi Murakami, Tatsuro Nakaji, Masahiro Nakamura, Rachakonda Sreekar, Legi Sam, Tomokazu Abe, Matthias Weiss, Katerina Sam

## Abstract

1. Birds, bats, and ants are recognized as significant arthropod predators. However, empirical studies reveal inconsistent trends in their relative roles in top-down control across strata. Here, we describe the differences between forest strata in the separate effects of birds, bats, and ants on arthropod communities and their cascading effects on plant damage.
2. We implemented a factorial design to exclude vertebrates and ants in both the canopy and understory. Additionally, we separately excluded birds and bats from the understory using diurnal and nocturnal exclosures. At the end of the experiments, we collected all arthropods and assessed herbivory damage.
3. Arthropods responded similarly to predator exclusion across forest strata, with a density increase of 81% on trees without vertebrates and 53% without both vertebrates and ants. Additionally, bird exclusion alone led to an 89% increase in arthropod density, while bat exclusion resulted in a 63% increase. Herbivory increased by 42% when vertebrates were excluded and by 35% when both vertebrates and ants were excluded. Bird exclusion alone increased herbivory damage by 28%, while the exclusion of bats showed a detectable but non-significant increase (by 22%). In contrast, ant exclusion had no significant effect on arthropod density or herbivory damage across strata.
4. Our results reveal that the effects of birds and bats on arthropod density and herbivory damage are similar between the forest canopy and understory in this temperate forest. In addition, ants were not found to be significant predators in our system. Furthermore, birds, bats, and ants appeared to exhibit antagonistic relationships in influencing arthropod density. These findings highlight, unprecedentedly, the equal importance of birds and bats in maintaining ecological balance across different strata of a temperate forest.

## Introduction

Arthropod herbivores play a critical role as primary consumers of leaf tissue in forest ecosystems (Coley, 1991; Coley & Barone, 1996). This may have various effects not only on individual plants but also on vegetation as a whole, such as plant growth and fitness (Garcia & Eubanks, 2019), species composition (Bagchi et al., 2014), and nutrient cycling (Belovsky & Slade, 2000; Chapman et al., 2003).

By feeding on arthropod herbivores, insectivorous predators indirectly increase plant biomass, creating what is commonly known as top-down control or a trophic cascade (Paine, 1966, 1980). However, the strength of top-down control by insectivorous predators varies due to factors such as prey availability (Garrett et al., 2022), predation rate (i.e., the consumption of prey by predators per unit of time; Thomine et al., 2020), and the magnitude of non-consumptive effects (i.e., alterations in prey behaviour in the presence of predators; Kollross et al., 2023). Given the complexity of food webs, the degree of top-down control is context-dependent, leading to variations even within a single forest and among different forest strata. Unfortunately, research on trophic cascades has so far predominantly focused on easily accessible forest understories (Denmead et al., 2017; Ocampo-Ariza et al., 2023), limiting our understanding of their full extent (e.g., forest canopy).

The impact of different predator groups on arthropod densities can vary, and when their effects overlap, it can become challenging to distinguish the individual contributions of each predator group (Mooney, 2007; Perfecto & Vandermeer, 1996; Richards & Coley, 2007; Sih et al., 1998). An obvious step to evaluating trophic cascades is thus to observe what happens when the abundance or community composition of predators is altered. To address this, a common experimental approach is the use of exclosure experiments (Maas et al., 2019), which exclude different predator groups from insect prey and foliage.

Vertebrate insectivores such as birds and bats often act as top predators of terrestrial arthropods (Böhm et al., 2011; Johnson et al., 2010; Karp & Daily, 2014; Maas et al., 2013; Mooney et al., 2010; Nyffeler et al., 2018). Previous research has generally emphasised the significant impact of insectivorous birds on arthropod communities (Bael et al., 2008; Mooney et al., 2010). However, it is important to note that most of these studies did not solely assess the effect of bird exclusion but rather considered the impact of both bird and bat exclusions combined (Greenberg et al., 2000; Holmes et al., 1979; Sam et al., 2023; Singer et al., 2017; Van Bael et al., 2003), with a few exceptions (Bouarakia et al., 2023; Cassano et al., 2016; Kalka & Kalko, 2006; Williams-Guillén et al., 2008). The classical approach of covering foliage with a net successfully excludes both predator groups but does not allow the predatory effects of each group to be assessed separately. As a result, the individual contributions of insectivorous birds and bats to top-down control remain poorly understood.

In addition to vertebrate insectivores, certain groups of arthropod insectivores, particularly ants, are also expected to have a significant role in trophic cascades. Although they have been extensively studied as natural enemies and biological control agents (Mestre et al., 2012; Philpott & Armbrecht, 2006; Rosumek et al., 2009; Schifani et al., 2020; Tobing & Kuswardani, 2018), their importance as key predators remains uncertain both in tropical and temperate forests (Pérez-Espona, 2021; Sanders & van Veen, 2011; Thurman et al., 2019) and various other habitats (Blaise et al., 2021; Bulgarini et al., 2021; Ohyama et al., 2020; Tuma et al., 2020). The ambiguity persists because only a limited number of studies have aimed to exclude ants whilst also considering vertebrate predators. Additionally, these studies often focus either on the forest understory (Denmead et al., 2017; Ocampo-Ariza et al., 2023) or the canopy (Singer et al., 2017). Findings have been mixed, with studies showing both negative (Denmead et al., 2017; Singer et al., 2017) and positive (Ocampo-Ariza et al., 2023) effects of ants on the abundance of mesopredators and herbivorous arthropods.

The combined predatory activity of multiple groups may have additive (Morrison & Lindell, 2012; Williams-Guillén et al., 2008), synergistic (Losey & Denno, 1998) or even antagonistic effects (Ferguson & Stiling, 1996; Mooney, 2007) on the control of arthropods. Compared to ants, insectivorous birds and bats primarily target larger arthropods (Philpott et al., 2004; Van Bael et al., 2003), and their diurnal and nocturnal foraging behaviour gives them access to distinct prey types. In such instances, their combined impact on arthropod communities equals the sum of the arthropods consumed by each group independently (i.e., additive effect). That considered, and acknowledging the distinct dietary preferences observed among ants, birds, and bats, we expect that they will evenly and strongly affect arthropod communities and that their effects will be additive **[H1]**.

It remains unclear whether, or under which conditions, predators indirectly affect plants (Mooney et al., 2010). This uncertainty emerges because predators can also function as intraguild predators, consuming predatory arthropods. This can have a negative effect on plants, and consequently, when predators simultaneously consume predatory arthropods and herbivores, their net effect on plants could be dependent on the balance between these two factors (Gras et al., 2016; Ocampo-Ariza et al., 2023). While the theory on trophic interactions predicts that the effect of vertebrate exclusion might be moderately counterbalanced by the release of spiders, carabid beetles and other predatory arthropods (Hölldobler & Wilson, 1990), the existence of such release in the absence of ants appears to be minimal, at least in tropical habitats (Gras et al., 2016). We can expect that the impact of birds and bats as intraguild predators on arthropods will have a relatively mild cascading effect on plant damage **[H2a]**. Conversely, ants with comparatively weaker intraguild predation abilities are likely to exhibit a more pronounced influence on plant damage **[H2b]**.

Forest canopies are crucial for the overall functioning of forest ecosystems (Ozanne et al., 2003). According to ecological theories, species interactions are more intense and species richness is higher in the warmer and more productive forest canopies than in the cooler understories (Basset et al., 2015; Janzen, 1970; Nakamura et al., 2017; Schemske et al., 2009). However, this is not always consistent with empirical findings. Various studies have reported an opposite or variable pattern of abundance and diversity of different arthropod taxa across tropical, subtropical, and even temperate forest canopies (Aikens et al., 2013; Basset et al., 2003; Compton et al., 2000; De Dijn, 2003; De Vries, 1988; DeVries et al., 1997; Haack et al., 2022; Hill et al., 1992; Intachat & Holloway, 2000; Larrivée & Buddle, 2009; Schulze et al., 2001; Ulyshen, 2011). Moreover, the effects of predators on plants through trophic cascades have seldom been investigated across the strata of temperate forests (Aikens et al., 2013; Böhm et al., 2011).

According to the optimal foraging theory, predators should allocate more time to foraging in areas with higher prey density to reduce search time (Balza et al., 2020; Emlen, 1966; MacArthur & Pianka, 1966; Piel et al., 2021). Additionally, the majority of birds in temperate forests are expected to be foraging strata generalists (Marra & Remsen Jr, 1997). Similarly, in the northern hemisphere, there is no clear stratification of bat species composition between the canopy and understory (Collins & Jones, 2009; Kalcounis et al., 1999; Plank et al., 2012; Zeus et al., 2017). In contrast, understory dominance is more pronounced among ants in temperate forests (Seifert, 2008). In light of these observations, we predict that the impact of predators on arthropods will be more pronounced in strata with higher arthropod densities, thereby attracting birds and bats **[H3a]**. Conversely, following our hypothesis [H2b], we expect a more pronounced effect of predators on plants in the forest understory, which is anticipated to be richer in ants in temperate forests **[H3b]**.

To address the aforementioned hypotheses, we individually excluded birds, bats and ants in a fully factorial design that allowed us to separate the effects of predators on the arthropod communities in the understory and canopy of a temperate forest in Japan. We complemented our research on the impact of predators on arthropods and plants by conducting surveys of the excluded predator communities.

## Materials and Methods

### Study site

We conducted our experiment in the Tomakomai experimental forest in Japan (42° 40 ’48.0”N 141° 35’ 24.0 ’’E, 50m a.s.l.). The study area covers a total of 2,720 hectares and has canopy crane access (Figure S1.1 in Appendix S1). It is situated on a hillside within the district, approximately 4 kilometres from the Pacific Ocean. The forest belongs to a cool, temperate zone and is composed of approximately 25% artificially planted conifers (e.g., *Picea glehnii* and *Abies sachalinensis*) and 75% young secondary deciduous forest, mainly occupied by broad-leaf trees dominated by oak (*Quercus crispula*), ash (*Fraxinus lanuginosa*), maple (*Acer mono, A. palmatum*) and elm (*Ulmus davidiana*), regenerated after typhoon damage (Wu et al., 2019). The temperature ranges from −22°C to 28 °C depending on seasonality. Annual precipitation ranges between 800 to 1,600 mm.

### Treatment preparation

We preselected eight plant species in the understory and seven in the canopy (Table S1.1) based on their abundance in the forest understory and canopy, and accessibility from the crane (Figure 1a). For each plant species, we identified suitable individuals - “saplings” (i.e., young trees, 1.5-3 m tall) in the understory and canopy “branches” (i.e., 1-1.5 m long branches of adult trees). The branches in the canopy had a comparable size and number of leaves to the saplings in the understory. They extended to heights ranging from 12 to 22 metres (Matsuo et al., 2022), resulting in a vertical separation of approximately 10.5 to 20.5 metres between the canopy and the understory.

**Figure 1.**
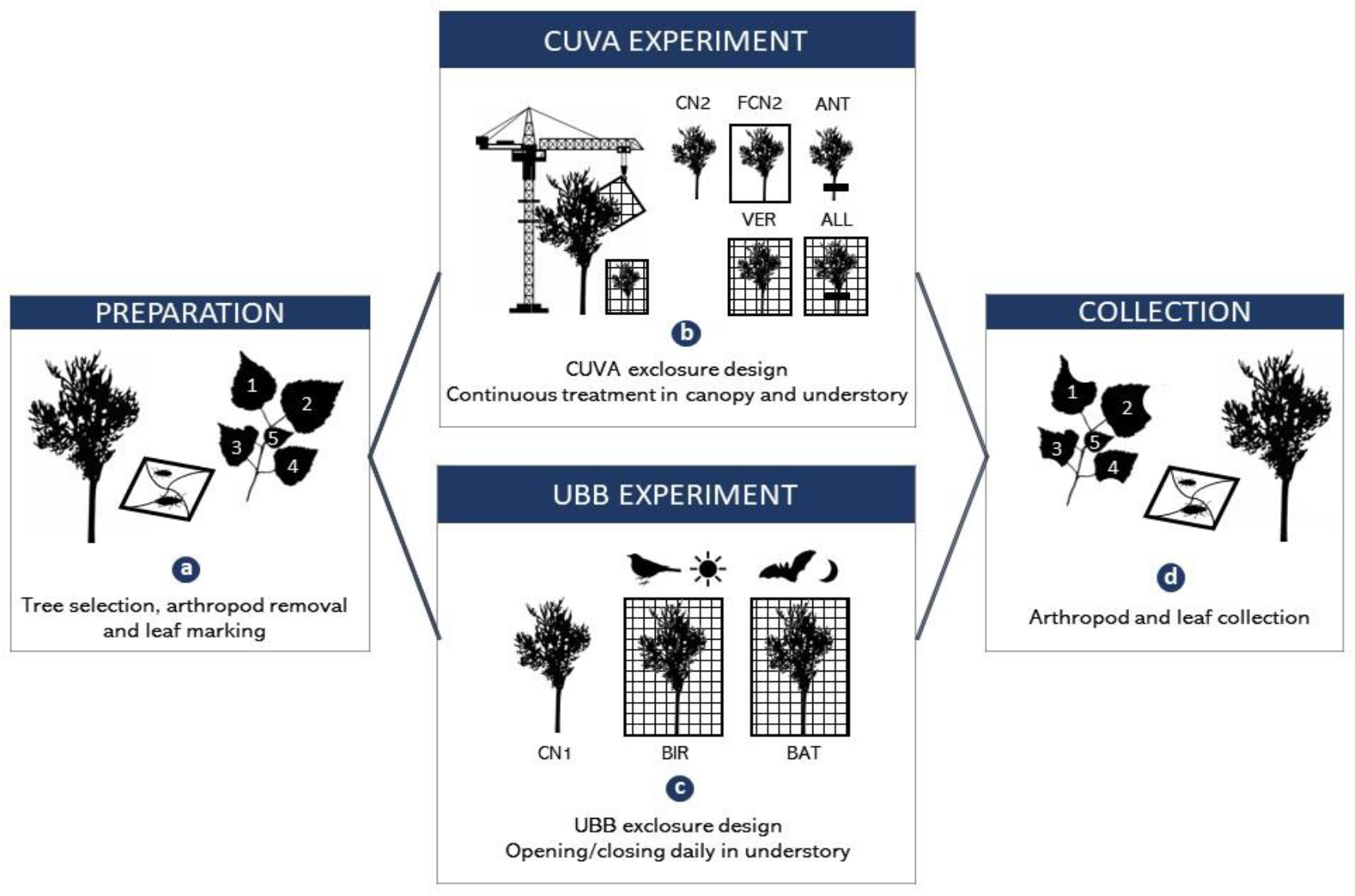
Schematic design of the experimental setup used in the study: (a) Pre-selection of individuals, removal of arthropods and assessment of preexisting herbivory on developing leaves to establish a baseline for the experiment; (b) Canopy - Understory Vertebrate and Ant exclusion (CUVA) experiment setup performed on branches in the canopy and on saplings in the understory; (c) Understory Bird and Bat exclusion (UBB) experiment, performed only on saplings in the understory; (d) Final arthropod collection and leaf herbivory survey at the end of each experiment. CN1 and CN2 = control treatments, FCN2 = frame-control treatment, ANT = ant exclosure, VER = exclosure of vertebrate predators, ALL = exclosure of vertebrates and ants, BIR - bird exclosure, BAT = bat exclosure.

We carried out two distinct experiments, each was conducted twice; first in 2018 and then replicated in 2019, using a distinct set of individuals each year. A total of 120 saplings were used each year for the Understory Bird and Bat exclusion (UBB) experiment, which was conducted exclusively in the understory. For the Canopy - Understory Vertebrate and Ant exclusion (CUVA) experiment, we used 84 branches and 160 saplings each year (Table S1.2 and S1.3). To establish a baseline at the beginning of the experiment, we removed all arthropods from the understory saplings and canopy branches. After the first leaves were fully developed, we randomly chose three small twigs from each sapling, once having at least ten leaves and no herbivory. Each twig was marked and had ten leaves numbered individually with a permanent marker. Then, we randomly assigned the individuals to a given experiment (UBB or CUVA) and a treatment.

### Canopy - understory vertebrate and ant exclusion (CUVA)

The CUVA experiment was conducted in both the canopy and understory (Figure S1.1), between May and July 2018 and 2019 (Table S1.4), until the end of the growing season which was determined by the leaf fall of *Prunus* species. The experiment lasted 65 ± 3 days. The experimental design consisted of setting up exclosures of vertebrates (VER), ants (ANT), and all predators combined (ALL). Each treatment was matched with a CUVA experiment control treatment (CN2), and a frame-control treatment (FCN2, Figure 1b). Each treatment was set on five individual saplings in the understory and three branches in the canopy for each of the eight/seven plant species in each year of collection (Table S1.2).

We constructed vertebrate exclosures (VER) using bamboo poles covered with agricultural transparent green netting, with a mesh size of 3 x 3 cm, which was comparable to mesh sizes used in other exclosure studies (e.g., Greenberg: 29×29 mm, Greenberg et al. 2000; Mols and Visser: 25×25 mm, Mols & Visser 2002; Van Bael: 20×20 mm, Van Bael et al. 2003). Each exclosure measured 2 x 2 x 2.5 m and had a total volume of 10 m3 in the understory, while in the canopy, they measured 1.5 x 1.5 x 1.5 m with a volume of 3.38 m3, enclosing an average of 1.63 m2 (± S.E. 0.06) of leaf area. The vertebrate exclosures were set permanently for the whole duration of the experiment in each year. The nets made firm contact with the ground in the lower part of the exclosures (Figure S1.2), and we securely fastened them to branches for canopy branches. Our observations confirmed that small insectivorous lizards and terrestrial mammals could access the exclosures, although we rarely observed them. We took special care to ensure that the foliage of the inner sapling did not touch the netting or cage construction, preventing flying vertebrates from accessing arthropods through the mesh. Importantly, the exclosure materials neither attracted arthropods nor caused damage to leaves or branches, and they did not reduce light exposure.

Ants (ANT) were excluded using a Tanglefoot sticky pest barrier (Philpott et al., 2004; Philpott et al., 2008). We applied the adhesive in a 10-15 cm wide stripe around the entire circumference at either the breast height of sapling trunks or the thickest section of canopy branches. Additionally, we removed all tall herbs and foliage in the surrounding area that could act as vegetation bridges, so the individual tree would not become accessible to ants after the treatment was set up.

To exclude all predators (ALL) we used a combination of the aforementioned methodology to exclude vertebrates and ants. Additionally, tangle glue was applied along any supportive ropes attached to the cage that could be used as a bridge for foraging ants.

The controls (CN2) were not enclosed by any constructions or protected by a tanglefoot barrier. The frame-controls (FCN2), on the other hand, were surrounded by identical bamboo constructions with dimensions 2 x 2 x 2.5 m (Figure S1.2). However, we did not surround these structures with agricultural nets. We used this treatment to investigate whether the construction had any unintended effects on the experiment, such as deterring vertebrate predators or attracting more mesopredators.

### Understory bird and bat exclusion (UBB)

The UBB experiment was conducted between May and June in 2018 and 2019 (i.e., the whole experiment was replicated twice) (Table S1.4). The exclosure experiment always lasted 30 ± 2 days. We set up exclosures for birds (BIR) and bats (BAT) with additional UBB experiment control saplings (CN1) (Figure 1c). Each of the three treatments was set on five individual saplings per plant species each year (Table S1.3). We exclusively conducted this experiment in the understory.

To exclude birds (BIR) or bats (BAT) separately, we used similar exclosure cages to those used for the VER treatments (Figure S1.2). The netting was pulled up to allow predators to access the sapling and down to exclude them. We moved the netting up or down ±30 min around sunrise and ± 30 min around sunset, ca. 4:15 AM and ca. 6:40 PM in mid-May, respectively, adjusted to the real sunrise and sunset daily. For BIR exclosures, we opened the exclosures during the night and closed them during the day and vice versa for BAT. As for CN2, individuals in CN1 were only marked.

### Collection of the experiments

The leaves marked at the beginning of the experiments were collected at the end of the experiments (Figure 1d). Individual leaves were scanned (EPSON, 600 dpi, colourful tiff format) within 12 hours of collection (Figure S1.2). To analyse insect herbivory, we first outlined any missing parts of leaves in Photoshop® using the protocol established by Sam et al. (2020). Then, we calculated the remaining area (a) and the full expected area of each leaf (b), in cm² using ImageJ version 1.47 (National Institute of Health, USA) in order to calculate the total area eaten by herbivores (c), (c = b - a) per leaf. We then calculated the proportion of leaf area loss as c/b. To determine the total leaf area of each individual sapling or branch, we calculated the mean leaf area based on the leaves collected for herbivory assessment. Then we multiplied it by the total leaf count in each sapling or branch using the mean value obtained from three independent estimations conducted by sampling technicians.

To survey the effect of our treatments at the end of each experiment, we first accessed the individuals and cut open the cages where needed. We lowered the crown of the individuals above a 1.5 x 1.5 m beating sheet, shook the foliage vigorously five times and quickly captured all arthropods that had fallen on the sheet (Figure S1.2). We then inspected the leaves for any concealed arthropods and took notes on any arthropods that escaped during the beating process. We stored the arthropods in vials filled with DNA grade 95% ethanol solution. All individual arthropods were later categorised into morpho-species, measured, and identified into their taxonomic order in the laboratory in the Czech Republic. Individuals were then assigned to one of four feeding guilds: predator, leaf chewer, sapsucker, or no relationship (i.e., arthropod with no consumptive effect on other arthropods or plants, NR).

The developmental stage of the arthropods was taken into account when assigning them to feeding guilds. For instance, adult Lepidoptera were classified as having “no relationship,” while their caterpillars were categorised as “chewers”. To calculate arthropod density, the number of arthropod individuals was determined per square metre of total leaf area of the individual sapling or branch. Closer identifications were done where needed to assign each individual to a given feeding guild.

### Predator survey

We used baits to survey both terrestrial (those that nest or forage on or in the leaf litter) and arboreal (those that forage or nest in the canopy) ants. The baits were exposed on eight randomly selected tree individuals of each of the eight focal plant species (i.e., 64 saplings in the understory and 64 branches in the canopy) during the second year of collection (i.e., 2019). We selected trees and branches different from those used in the exclosure experiment for the ant survey to avoid the disturbance of the experiment. Yet, the trees used for the ant survey were growing in the same plot but randomly scattered among the individuals used for the exclosures and were of a similar size and amount of foliage. Two types of baits (each roughly 2-3 cm^3^) were used: 1) tuna chunks in vegetable oil (Giana®) and 2) cotton balls soaked in sugar paste and wrapped in a piece of gauze. Baits were set out at each selected sapling or canopy branch, attached with a string and separated by at least 20 cm, and an alternating position of the bait was used (i.e., lower or higher on the trunk of saplings, and either closer or further from the trunk on canopy branches, or altering positions on a fork-shaped branch if available). We checked the baits after 4 hours of exposure and visually morphotyped and counted any ants feeding on them. Up to five individuals of each morphotype crawling on the bait were collected and put into 2 ml vials filled with DNA-grade ethanol (99%). During sampling, information about the time, weather conditions, bait position, ant morphotype and the abundance per morphotype was recorded on datasheets. We later identified ants in the laboratory using the species level key (Ichinose, 1990; Imai et al., 2003) based on records of existing ant species at the study site (Ichinose, 1990). While collecting the baits, the surrounding branches were examined for additional ants that may have been feeding on the baits prior to inspection.

We used point counts and Song Meter recordings to assess bird communities. Point counts were carried out at 16 points regularly spaced along a 2,350-m transect at the study site; successive points were 150 ± 5 m apart to avoid overlap and up to 120 m apart in elevation. All birds seen or heard within a fixed radius of 0–50 m (estimated or measured by a laser rangefinder) were recorded (Sam et al., 2019), and the height of the individual above ground was noted. We started surveys 15 minutes before sunrise, each count lasted 15 minutes so that all 16 points were surveyed before 11:00 (i.e., such a survey on all 16 points represents one replication in time) (Sam et al., 2019) during the second year of collection (i.e., 2019). All points were surveyed equally, and the survey was replicated fifteen times (i.e., in 15 days). A Song Meter SM3BAT (Wildlife Acoustics Inc.) with one external acoustic (SMM_A1, Wildlife Acoustic) and one ultrasonic (SMM_U1, Wildlife Acoustic) microphone was set to record the first 10 minutes of every 30 minutes (10 min recording, 20 min sleeping) from 3:00 AM to 9:00 PM. The Song Meter was set up for 15 days in the forest canopy (17m above the ground) during the first month of the experiment and for 15 days in the forest understory (1.5m above the ground) during the second month of the first year of collection (i.e., 2018). We used manual identification of the bird calls from the recordings and determined species richness and relative abundances of birds in the understory and canopy.

Similarly, we estimated bat communities using the same Song Meter. The Song Meter was set for 14 days in the forest canopy and 14 days in the forest understory during the second year of collection (i.e., 2019) using the same parameters as the bird survey. The recordings were divided into five-second files and analysed by opening each WAV file in Kaleidoscope Pro Software (Wildlife Acoustics Inc.) and manually inspecting the spectrograms for bat echolocation pulses. The sampling rate was 192 kHz. Echolocation call types were recognised from the recordings and attributed to a bat species (when possible) based on literature (Fukui et al. 2004). We measured bat activity from the Song Meter recordings as a proxy of abundance. We defined a ‘bat pass’ as a sequence with at least two recognisable echolocation pulses per species emitted by a flying bat within a 5-second sound file (Kerbiriou et al., 2019). Bat activity was quantified as the number of bat passes recorded for each species. Later on, we used the Handbook of the Mammals of the World (Zachos, 2020) and Handbook of the Birds of the World (Del Hoyo et al., 1996) to obtain the body weight and feeding guild of each bird or bat species to determine the biomass of insectivorous predators at each forest strata.

### Statistical analysis

We first built linear mixed-effect models using the package “lme4” (Bates et al., 2015), to test the effect of treatment (factor of 4 levels), strata (factor of 2 levels) and their interaction on log-transformed total arthropod densities (number of individuals per cm² of foliage) and arthropod densities partitioned into four feeding guilds: chewers, mesopredators, sapsuckers and no relationship (hereafter referred to as NR) for the CUVA experiment. All the models additionally contained the sampling year (factor of 2 levels) as a fixed effect and individual trees (factors of 361 levels) and plant species (factor of 8 levels) as random effects. We also considered strata both as a fixed effect and as a random slope for the plant species (referred to as Plant species: Strata in Table 3) to account for species differences between strata. Then, we ran generalised linear mixed-effect models using the package ‘glmmTMB’ (Brooks et al., 2017), using a beta error distribution and the same predictors as above to model herbivory damage (the proportion of the leaf area lost per branch). All the models contained the sampling year as a fixed effect and individual trees, branches (factor of 487 levels) and plant species as random effects. We again considered strata both as a fixed effect and as a random slope for the plant species. To select the best models, we used the AICctab function from the ‘bbmle’ package (Bolker & Bolker, 2017), which computes the corrected Akaike information criterion of all our models (Table S2.1, S2.2). For each best model, we obtained estimated marginal means (= emmeans) and comparisons among all variable levels (Table S2.3, S2.4, S2.5, S2.6, S2.7), using the ‘emmeans’ package (Lenth, 2018). We controlled the model’s quality and fit with the ‘performance’ package (Lüdecke et al., 2021).

We then constructed linear mixed-effect models to test the effect of treatment (factor of 3 levels) on log-transformed total arthropod densities and arthropod densities partitioned in four feeding guilds: chewers, predators, sapsuckers and NR for the UBB experiment. All the models contained the sampling year as a fixed effect and plant species (factor of 8 levels) as a random effect. Then, generalised linear mixed-effect models using a beta error distribution were used to determine the effect of treatment on herbivory damage. All the models contained the sampling year as a fixed effect and plant species and individual trees (factor of 240 levels) as random effects. The best models were selected following the same method as for the CUVA experiment (Table S2.8, S2.9), as well as for their estimated marginal means (Table S2.10).

We used Wilcoxon rank sum tests to compare total arthropod densities, predator, chewer, sapsucker and NR densities, and herbivory between canopy and understory control treatments (CN2) as well as total arthropod and mesopredator densities between control (CN2) and frame-control (FCN2) in the understory. All analyses were performed using R Statistical Software (v4.3.1; R Core Team, 2020).

## Results

### Arthropod density and herbivory damage after the CUVA experiment

In total, we collected 10,649 arthropods from 488 individual branches and saplings across 9 plant species (i.e., two years combined). We found that the overall arthropod densities, as well as predatory arthropod densities, on control individuals, were significantly lower in the canopy than in the understory (W = 2245, P = 0.002 and W = 2539, P < 0.001) (Figure S3.1), whereas the arthropod chewer, sapsucker and NR densities on controls did not vary significantly between forest strata (see Table 1 for more details).

**Table 1:** Mean arthropod density per square metre (i.e., total and split into feeding guilds) and herbivory damage (%) found in the control individuals in the canopy and understory (± standard error). Significant comparisons between the canopy and understory are marked as follows: * P ≤ 0.05, ** P ≤ 0.01, *** P ≤ 0.001.

| Strata | Total arthropods ** | Chewers | Predators *** | Sapsuckers | NR | Herbivory (%) *** |
| --- | --- | --- | --- | --- | --- | --- |
| Canopy | 22.2 ± 3.9 | 2.8 ± 0.5 | 5.0 ± 1.0 | 2.9 ± 0.6 | 11.4 ± 3.2 | 5.1 ± 0.5 |
| Understory | 46.1 ± 6.0 | 13.2 ± 3.9 | 24.3 ± 4.3 | 1.9 ± 0.5 | 6.6 ± 1.4 | 8.5 ± 0.3 |

Ant exclusion was relatively effective, we only collected a total of 49 ants in ALL and ANT understory treatments (which excluded ants) and 3 ants in the canopy. In contrast, 140 and 100 ants were found in VER and CN2 understory treatments respectively, accounting for approximately 7% of all collected arthropods, while only 4 ants were recorded in VER and CN2 in total in the canopy (Table S3.1).

In both the canopy and understory, only VER and ALL exclusions led to a significant increase in arthropod density (Figure 2a). Arthropod density increased by 82 % in VER and 53 % in ALL compared to the controls (z = −5.086, P < 0.001; z = 3.631, P = 0.001, respectively) (Table 2) in both the canopy and understory. The effect of ANT exclusion did not differ significantly from the control in both the canopy and the understory (Table 2 and Table S2.11).

**Figure 2:**
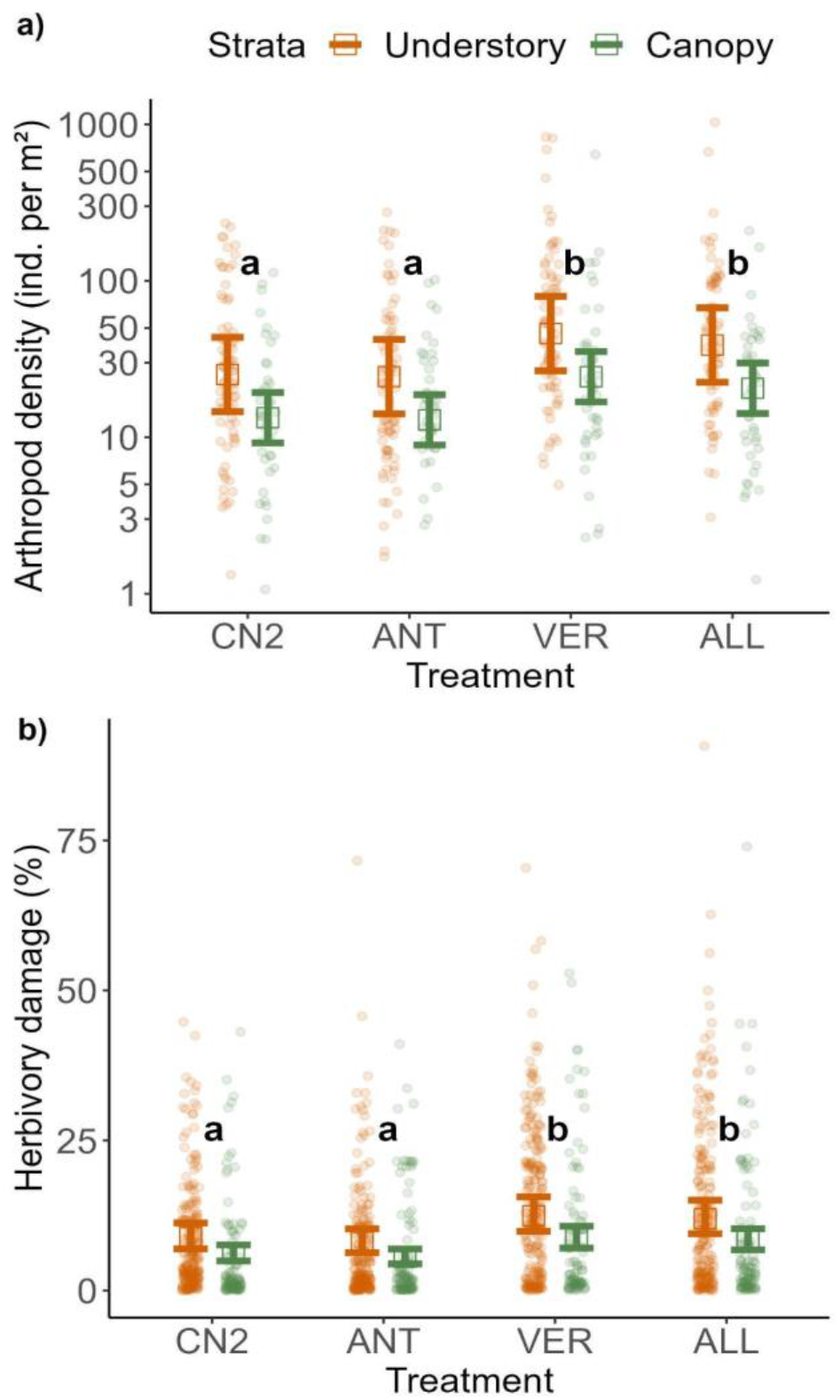
Total densities of all arthropods per square metre of foliage (a) and herbivory damage (%) (b) on surveyed saplings and branches of the CUVA (Canopy and Understory Vertebrate and Ant exclosure) experiment. Each individual data point represents (a) the density of arthropods or (b) the percentage of herbivory damage on either a canopy branch (green) or sapling individual in the understory (orange). The y-axis of (a) is on a log scale. Square and whiskers mark estimated marginal means and standard errors of the most parsimonious model. Significant pairwise comparisons between predictors were tested by Tukey post hoc tests and are indicated with letters (note that the results are the same for canopy and understory). CN2 = control treatment, ANT = ant exclosure, VER = vertebrate exclosure, ALL = all predator exclosure.

**Table 2:**
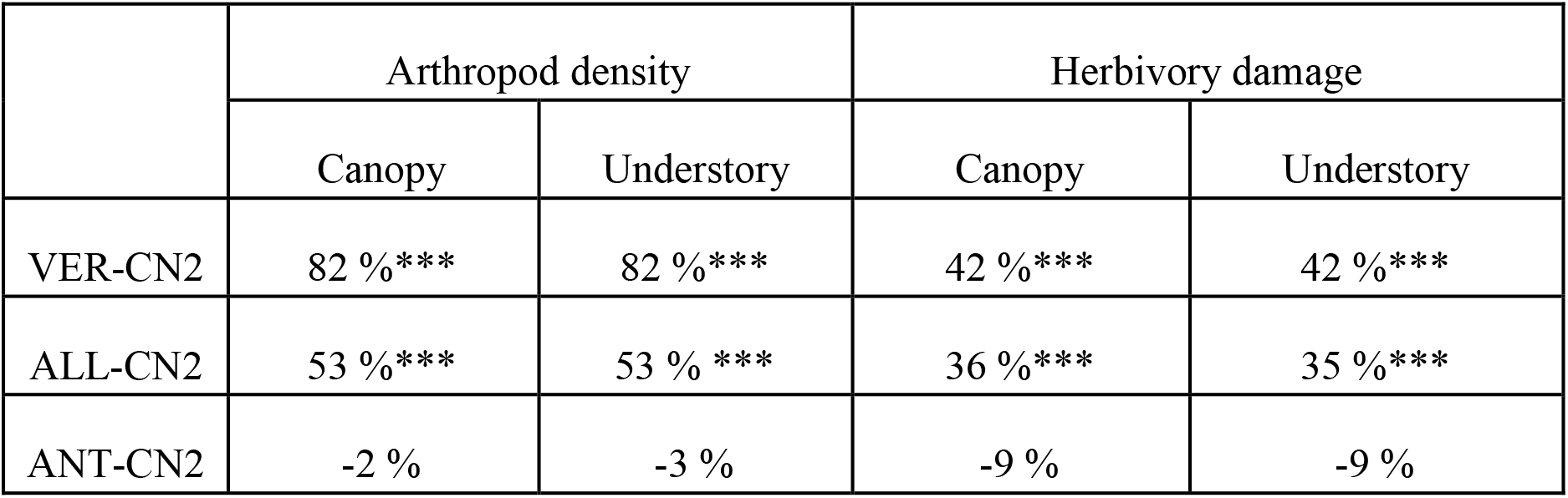
The percentage increase or decrease of arthropod density and herbivory damage between VER, ALL, and ANT treatments compared to CN2 (controls). The significance is marked as follows: * P ≤ 0.05, ** P ≤ 0.01, *** P ≤ 0.001.

|  | Arthropod density |  | Herbivory damage |  |
| --- | --- | --- | --- | --- |
|  | Canopy | Understory | Canopy | Understory |
| VER-CN2 | 82 % *** | 82 % *** | 42 % *** | 42 % *** |
| ALL-CN2 | 53 % *** | 53 % *** | 36 % *** | 35 % *** |
| ANT-CN2 | -2 % | -3 % | -9 % | -9 % |

Among the control treatment individuals, herbivory damage was significantly higher in the understory than in the canopy (W = 1290370, P < 0.001) with a 60 % increase in herbivory in the understory (Table 1).

In both the canopy and understory, only VER and ALL exclusions led to a significant increase in herbivory damage. In the canopy, herbivory increased by 42 % in VER and 36 % in ALL (Table 2) compared to the controls (z = −5.386, P < 0.001; z = 4.726, P < 0.001, respectively) (Figure 2b). In the understory, herbivory increased by 42 % and 35 % (Table 2) in comparison to the controls (P < 0.001) (Figure 2b). The effect of ANT exclusion did not differ significantly from the controls in both the canopy and understory (Table 2).

### Arthropod density and herbivory damage after the UBB experiment

In total, we collected 2,544 arthropods from 240 saplings across 8 plant species at the end of the UBB experiments (i.e., two years combined). The exclusion of birds led to a significant increase in mean arthropod density by 89 % in comparison to the control trees (z = 3.789, P < 0.001). Similarly, in the absence of bats, arthropod density increased significantly by 63 % in comparison to the control saplings (z = 2.876, P = 0.012) (Figure 3a).

**Figure 3:**
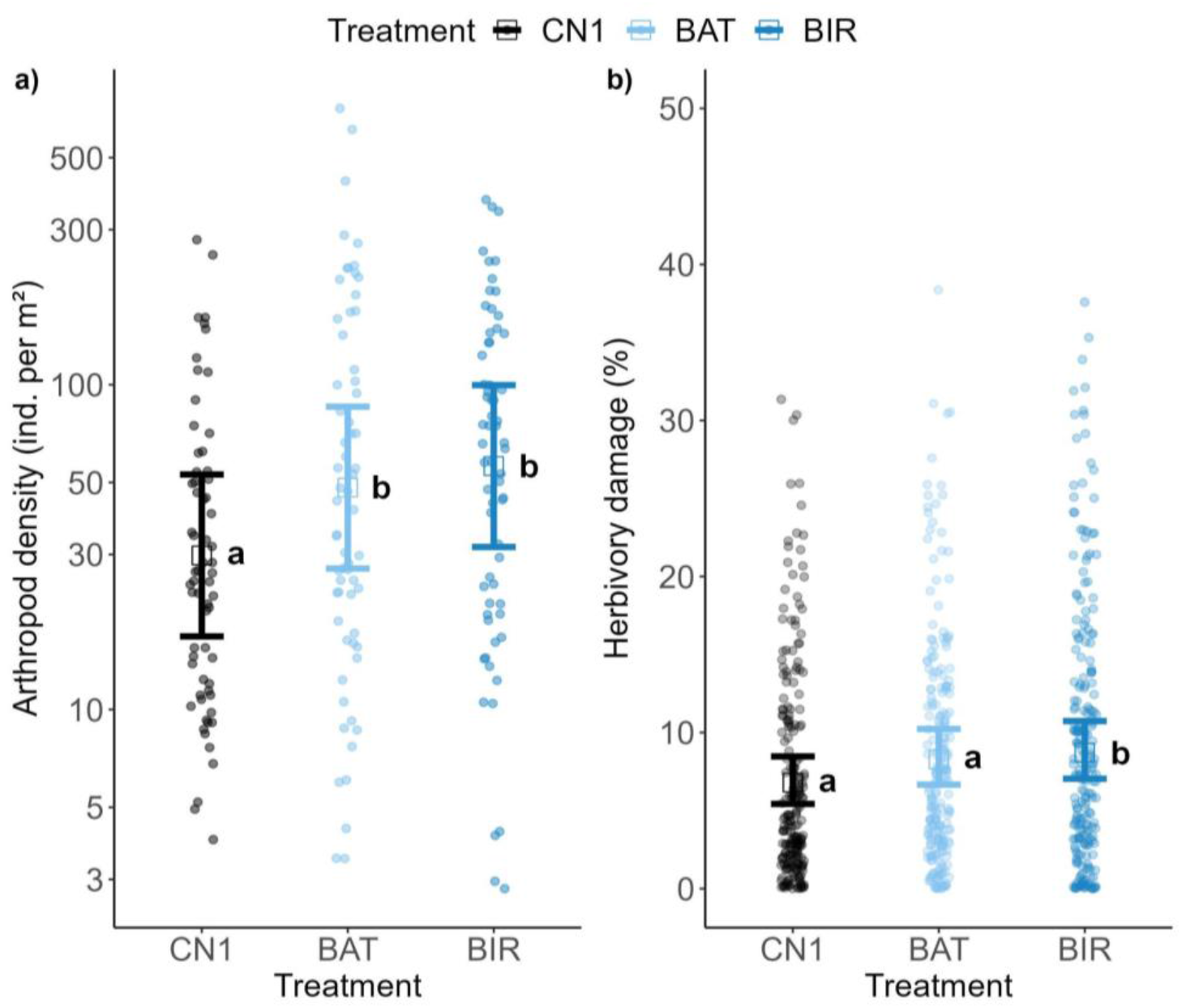
Effects of the exclusion of birds and bats on the total densities of all arthropods per square metre of foliage (a) and on the herbivory damage of individual saplings of the UBB (Understory Bird and Bat exclusion) experiment. Individual data point represents a density of arthropods (a) or herbivory damage (b) on an individual sapling. The y-axis of (a) is on a log scale. Square and whiskers mark estimated marginal means and standard errors of the most parsimonious model. Significant pairwise comparisons between predictors were tested by Tuckey post hoc tests and are indicated with letters. Note that four extreme values of herbivory have been removed from the BIR raw data for visualisation purposes (b). CN1 = control treatment, BAT = bat exclosure, BIR = bird exclosure.

However, only the exclusion of birds led to significantly increased mean herbivory damage (z = 2.827, P = 0.013) by 28% in comparison to the controls (Figure 3b). The effect of bat exclusion was detectable (+ 22% in comparison to the control) but non-significant (z = 2.221, P = 0.067).

### Arthropod feeding guilds

During the CUVA experiment, sapsuckers were the most abundant feeding guild (5403 individuals), found on 48% of saplings and branches, followed by no relationship arthropods (2231, present on 73%), predators (1865, present on 84%), and leaf chewers (1150, present on 58%). In the UBB experiment, predators (894, present on 89% of saplings) and sapsuckers (662, present on 68% of saplings) were dominant, followed by leaf chewers (526, present on 72%) and NR (462, present on 57%).

Across the CUVA experiment, only the sapsucker (z = 3.531, P = 0.001; z = 4.547, P < 0.001) and NR (z = 2.529, P = 0.032; z = 2.575, P = 0.028) arthropods exhibited significant increases in their densities, with increments of 297% and 161% after the exclusion of vertebrates (VER) and 504% and 166% after the exclusion of all predators (ALL), respectively (Figure 4a). Additionally, the exclusion of ants (ANT) resulted in a significant 64% reduction in the densities of predatory arthropods (z = −2.442, P = 0.041). None of the exclusions had a significant impact on chewer densities (Table S3.2). It is important to note that the strata did not exhibit significant interactions with the treatments for any of the arthropod densities (Table S2.2); therefore, they were not considered in these results.

**Figure 4:**
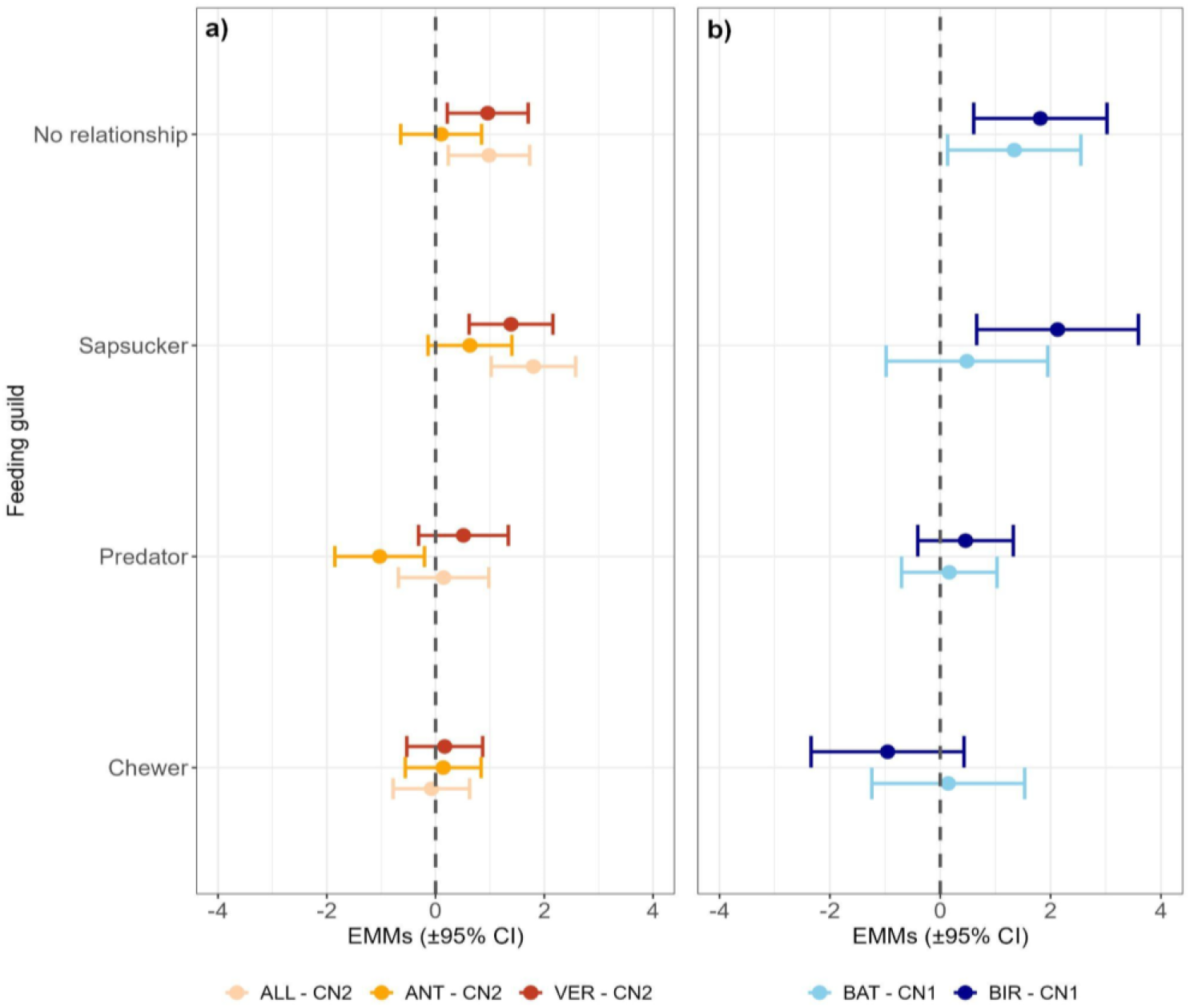
Caterpillar plot showing how the feeding guilds (predatory arthropods, chewing herbivores, sapsucking herbivores and NR) respond to (a) all (ALL, beige), ant (ANT, orange) and vertebrate (VER, red) exclusion treatments in the CUVA (Canopy and Understory Vertebrate and Ant exclosure) experiment using canopy and understory data combined and to (b) bat (BAT, light blue) and bird (BIR, dark blue) exclusion treatments in the UBB (Understory Bird and Bat exclusion) experiment (understory only). The X-axis shows the estimated marginal means (EMMs) of each treatment against the control (dashed line = no change from control) with a 95% confidence interval of the most parsimonious model including the variable treatment. When the confidence interval is strictly above or below the dashed line, the effect is significant. CN2 = control treatment for the CUVA experiment. CN1 = control treatment for the UBB experiment.

At the end of the UBB experiment, the NR (z = 2.17, P = 0.058; z = 2.94, P = 0.007) significantly increased their densities after excluding bats and birds independently by 285 and 517 % respectively. In addition, the sapsucker arthropods significantly increased their densities by 741 % after the removal of birds (z = 2.84, P = 0.009). Neither the chewers nor the predatory arthropods significantly changed their densities after the exclusion of bats and birds (Figure 4b).

We did not find significant differences in the densities of total arthropods (W = 1491, P = 0.545) and predatory arthropods (W = 1626, P = 0.887) between controls (CN2) and frame-controls (FCN2) (Figure S3.2, S3.3, respectively).

### Vertebrate and arthropod predators

Overall, we identified a total of 22 insectivorous bird species comprising 1,167 bird calls using point counts and recordings (i.e., two methods combined), as well as 5 insectivorous bat species consisting of 79 bat passes from recordings and 7 ant species from 908 individuals caught on baits (Table S3.4). The surveys conducted across different strata revealed that insectivorous birds were 139 % more abundant in the canopy than in the understory whereas insectivorous bat activity and ant abundance were 238 and 1,264 % greater in the understory than in the canopy (Figure 5a). For insectivorous birds, species richness was similar within strata, while a greater number of ants and bats (i.e., 40 and 33 % more species, respectively) were found in the understory (Figure 5b). In addition, bird biomass was 90% greater in the canopy than in the understory, whereas bat biomass was 258% greater in the understory than in the canopy (Figure 5c).

**Figure 5:**
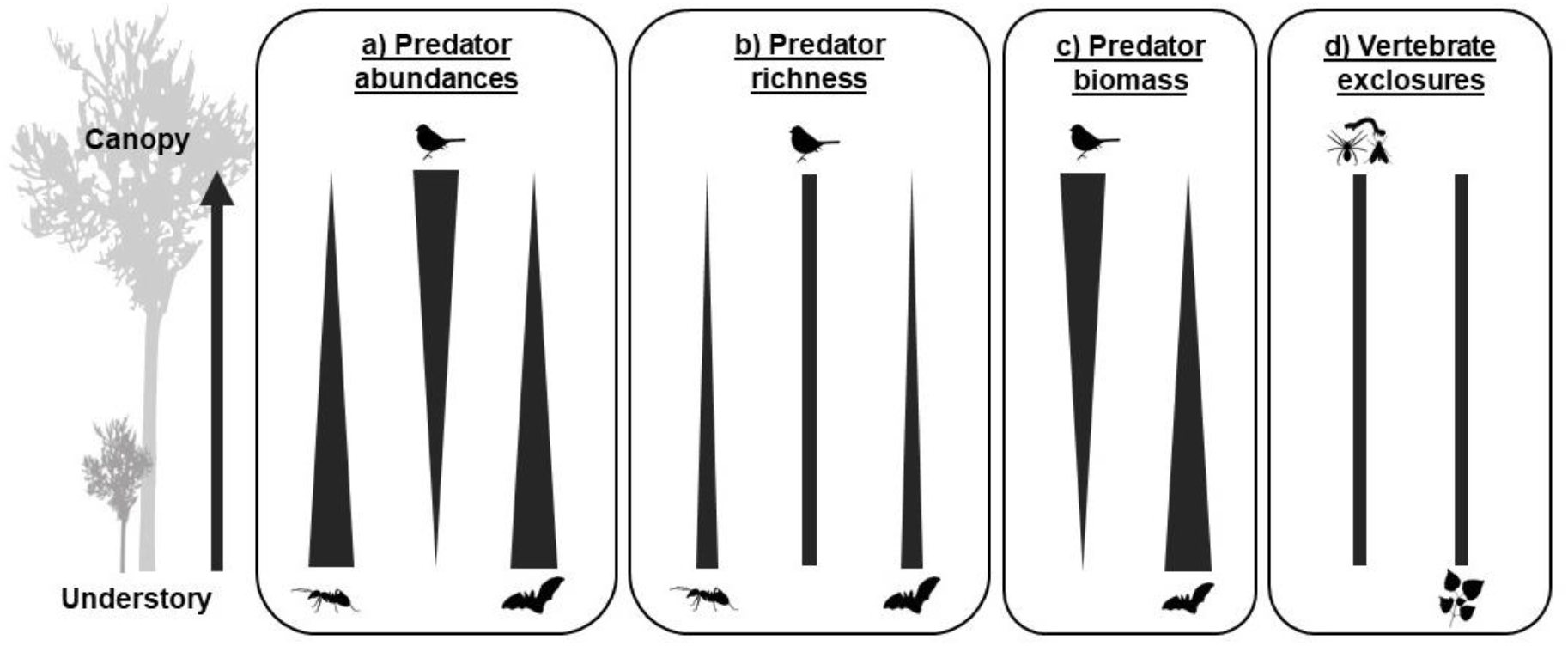
Simple comparisons between canopy and understory levels for (a) insectivorous predator abundances (ants, birds, bats) (b) richness (ants, birds, bats) (c) biomass (birds and bats) (d) CUVA (Canopy and Understory Vertebrate and Ant exclosure) experiment results (effect on arthropod densities and herbivory damage in VER). Note that the size of the arrow depends on the magnitude of the difference between the canopy and the understory.

## Discussion

Vertebrate predators (i.e., birds and bats together), but not ants, played a crucial role in preventing arthropod outbreaks and indirectly protecting plants from herbivory damage. Our results reveal that their effect was similarly strong in both the forest canopy and understory. Specifically, the absence of vertebrates led to nearly double the density of arthropods, indirectly increasing the herbivory damage by almost half, regardless of the strata. Additionally, the individual effects of the bird, bat and ant exclusions demonstrated an antagonistic rather than an additive impact on arthropod density. In contrast to vertebrate predators, ants did not significantly impact either arthropod communities (except for mesopredators) or herbivory damage in any forest strata. Our study underscores the significance of vertebrate predator presence in maintaining the equilibrium of trophic cascades in both the canopy and understory of temperate ecosystems. These findings contribute to our understanding of the role of stratification in trophic cascades within temperate regions, an area that has been relatively unexplored in previous research (Aikens et al., 2013; Böhm et al., 2011).

To provide context, the density of arthropods observed on control individuals revealed a consistent trend: there were more arthropods per leaf area in the understory compared to the canopy, irrespective of their trophic roles, except for the sapsuckers. This trend was particularly pronounced among predatory arthropods in contrast to chewers, aligning with the established notion that vertical stratification of herbivores in temperate forests is relatively weak (Basset, 2003). Higher abundances or densities of arthropods in the understory have been attributed to various factors, including the greater stability of the microclimate closer to the ground (Parker et al., 1995), dispersal limitation after emergence (Brown, 1997) and the distribution of quality food resources (Basset, 2003). In line with the trend in arthropod distribution, we observed a 60 % increase in herbivory damage on understory control saplings when compared to canopy control branches. Additionally, mean herbivory damage was approximately 8% in our study, which closely matches the findings from previous studies in temperate forests, which typically report herbivory damage levels around 5-10% (Gossner et al., 2014; Reynolds & Crossley, 1997; Wang et al., 2016).

In contrast to our first hypothesis [H1], which postulated that predators would have a strong and additive effect on arthropod communities, our results indicate that vertebrate predators (VER) and ants (ANT) exhibit antagonistic effects when compared to the combination of vertebrate predators with ants (ALL). Similarly, bats (BAT) and birds (BIR) show antagonistic effects when compared to vertebrate predators (VER). This could imply that despite their differing activity periods, birds and bats may be partially competing for the same prey resources. Indeed, birds and bats both display a preference for large prey (Philpott et al., 2004; Sam et al., 2023; Sivault et al., 2023; Van Bael et al., 2003), potentially leading to a constraint in prey availability, thereby diminishing their collective effect. On the other hand, it is possible that the differing collection times of the two experiments (i.e., a one-month difference) might account for the observed pattern (Figure 6). We collected arthropods from the VER exclosures later in the season, which could lead to differences in prey availability and size.

**Figure 6:**
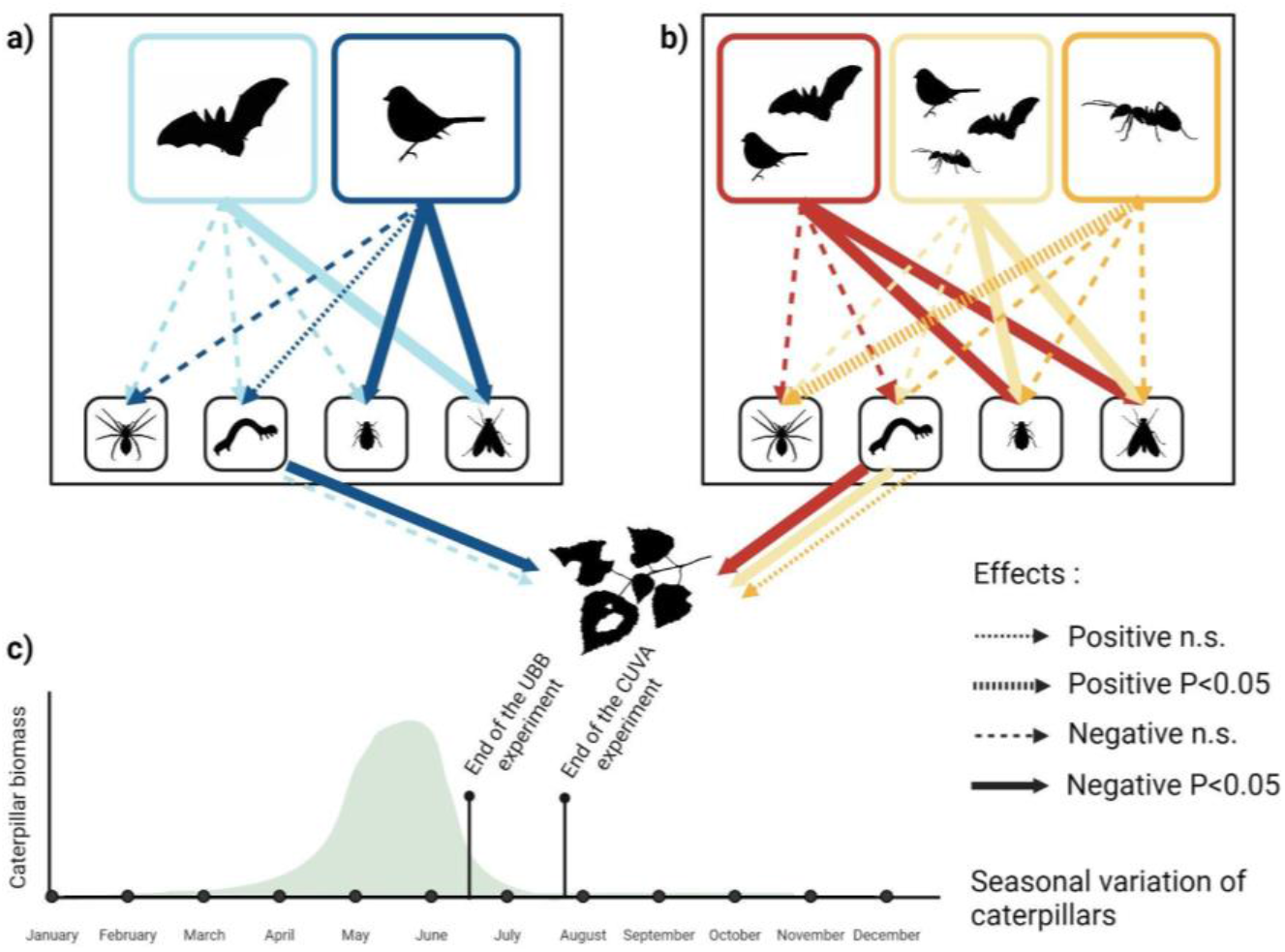
Distinct effects of bats and birds (UBB experiment - Understory Bird and Bat exclosure) (a), combined (VER), combined with ants (ALL) or predatory ants alone (ANT) (CUVA experiment - Canopy and Understory Vertebrate and Ant exclosure) (b) on mesopredators (spider), chewers (caterpillar), sapsuckers (aphid), NR (fly) densities and herbivory damage (leaf) (canopy and understory combined). Effects were assessed through generalised linear mixed models (GLMMs) (Table S2.2 and Table S2.9). The generic seasonal variation of caterpillar biomass in Japan, accompanied by the time frame of our experiments, is depicted at the bottom of the plot, following the results of Verboven et al. (2001), Murakami (2002), and Sayama et al. (2012) (c).

Furthermore, during the UBB and CUVA experiments, only BIR, BAT, VER, and ALL treatments significantly prevented arthropod outbreaks in both the canopy and understory, whereas the ANT treatment did not. These results contrast with previous studies in temperate forests where ants were shown to reduce the abundance of nearby insects (Gras et al., 2016; Sanders & van Veen, 2011). Looking at the ant species found during our surveys (Table S3.4), all of them are generalists, suggesting that ants were not a particularly important mesopredator in our system. In fact, their mutualism with sap-sucking insects may have been more relevant than their potential role as mesopredators in our study (Offenberg, 2001). Furthermore, the relative abundance of ants (ca. 7% of all arthropods) makes them important prey items for vertebrates.

Both the results of the UBB and CUVA experiments also showed that the removal of vertebrates mainly affected the densities of NR arthropods and sapsuckers, but not chewers and mesopredators (Figure 6). The lack of effects on chewers appears to be related to the difference in time frame between the two experiments. Due to the necessary duration constraints of the CUVA experiment required for the effective accumulation of predator and herbivore effects, the arthropod collection occurred after the peak of caterpillar abundances observed previously in similar Japanese forests (Murakami, 2002; Sayama et al., 2012; Verboven et al., 2001). Therefore, we did not collect many chewers at the end of the CUVA and UBB experiments, diminishing the overall effects of treatments on chewers. Surprisingly, ant exclusion led to a significant reduction in mesopredator densities. This could be because ants were classified as mesopredators in our study, and their removal may impact the overall mesopredator pattern. Additionally, ants can serve as an important food source for other predatory arthropods through myrmecophagy, such as spiders (Aranea), bugs (Heteroptera; Brandt & Mahsberg, 2002), net-winged insects (Neuroptera), or flies (Diptera; Aceves-Aparicio et al., 2022; Wilson, 2000).

In treatments where birds were absent (VER, ALL, and BIR exclosures), we observed an increase in sapsucker densities. This was unexpected, as we did not anticipate birds to feed abundantly on sapsuckers. Their small size and sessility likely make them inconspicuous to vertebrate insectivores which is consistent with recent findings in tropical areas (Ferreira et al., 2023; Ocampo-Ariza et al., 2023). It, therefore, seems likely that the increase in their densities observed here is due to an indirect interaction with vertebrate insectivores, since some insectivorous birds (e.g., *Parus minor, Sitta europaea*) found abundantly in Tomakomai forest consume other taxa, such as mesopredators (ants, wasps or spiders), and mutualists (ants) (Eguchi, 1980; Wesołowski et al., 2019), that positively affect the sapsuckers by relieving predation pressure.

Regarding mesopredator densities, vertebrates (VER) did not act as intraguild predators significantly in the CUVA experiment, therefore contrary to [H2a], their net effect on herbivory damage appeared significant (Figure 6). The 42 % increase in herbivory in the vertebrate exclosures, compared to the controls, was comparable to the change in damage found after vertebrate predator exclusion in a German canopy (ca. 23 to 44%; Böhm et al., 2011), but greater than in other temperate canopies (ca. 0-15 %; Barber & Marquis, 2009; Beilke & O’Keefe, 2023; Lichtenberg & Lichtenberg, 2002) and understories (ca. 0-18%; Barber & Marquis, 2009; Dekeukeleire et al., 2019; Maguire et al., 2015).

Bird exclosures led to an indirect increase in herbivory damage, whereas bat exclosures had a weak impact within the UBB experiment. Moreover, the effects of birds and bats on herbivory seemed to be additive. It is possible that the effect on herbivory damage was more detectable than changes in arthropod densities, as herbivory accumulated throughout the experiments, whereas arthropod collection was confined to a specific day. Consequently, the magnitudes of these effects exist on distinct scales, and they are markedly biased. Contrary to the second part of the second hypothesis [H2b] but following their weak effect on arthropods, the effect of ants on herbivory damage was not detectable (Figure 6), when combined with vertebrates, the effect remains significant but to a lesser extent.

In contrast to the third hypothesis that the effect of predators will be more pronounced in the strata with higher arthropod densities [H3a], our observations revealed that vertebrate predators had a similar impact on arthropod densities both in the forest canopy and understory. This result aligns with other studies from temperate forests where vertebrate predation pressure on arthropods did not differ between vertical strata (Aikens et al., 2013; Boege & Marquis, 2006). Yet, it contradicted the optimal foraging theory, as we observed lower arthropod densities in the canopy than in the understory. Our predator surveys revealed that the abundance of insectivorous birds was 140% higher in the canopy than in the understory. In contrast, bats and ants were found to be the most abundant and rich in the understory. Therefore, the similar impacts on arthropods observed across strata may be attributed to the balanced predation pressure, with an increased presence of birds in the canopy and greater abundances of bats and ants in the understory. This stratification of bats followed previous observations in the Tomakomai experimental forest, where they exhibit some degree of niche partitioning (Fukui et al., 2004); however, it is important to note that the bat sampling effort is much lower than that for birds in this study, as such, these results warrant a nuanced interpretation.

Our surveys confirmed the partitioning of ant species among distinct forest strata, with the majority of ant species being located in the understory (Seifert, 2008). Despite a relatively modest count of ant morphospecies (7 morphospecies in the forest understory and 5 morphospecies in the forest canopy), the use of ant baits indicated a considerably higher ant abundance in the forest understory. This pattern aligns with a comprehensive study conducted by Floren et al. (2014), which identified that arboreal ants exhibited only 12 % of the species found in temperate sites. This could be attributed to the ability of ants in tropical forests to construct additional nesting forms, such as epiphytic nests. These nesting structures are frequently absent in temperate forests, primarily due to differences in climatic conditions (Blüthgen & Feldhaar, 2010; Liefke et al., 1998), which limit the opportunity for many ant species to adapt to canopy conditions (Floren et al., 2014).

Our results contradicted the second part of our third hypothesis [H3b], where we anticipated a stronger effect of predators in the forest understory. We found that the herbivory damage, resulting from trophic cascades between predators and plants, was consistent between the forest canopy and understory, likely independent of ant activity. As previously noted, the effect of ants on arthropod communities, and consequently their impact on herbivory, appears to be relatively minor within the studied temperate forest. These results follow the same pattern as arthropod density, suggesting that predation pressure on phytophagous insects may also be consistent within strata. However, we failed to detect significant changes in the density of chewers in our study, likely due to the phenology of caterpillars (Figure 6). To our knowledge, there is limited research on herbivory levels in the understory and canopy of temperate forests, (but see Gossner et al., 2014; Reynolds & Crossley, 1997; Wang et al., 2016 for the herbivory levels in temperate understories). Therefore, further research is necessary to determine if this observed pattern is consistent across other temperate forests.

## Conclusion

Our study demonstrates that birds and bats, but not ants, play a crucial role in reducing arthropods in both the temperate forest canopy and understory, leading to a significant decrease in plant damage. Given that insectivorous bats and birds are present in numerous terrestrial ecosystems (Maas et al., 2013), the importance of their predation likely extends to many other areas. In addition, being threatened by various factors worldwide such as habitat loss, hunting, and the impacts of climate change (Benitez-Lopez et al., 2017; Donald et al., 2001; Stephens et al., 2016), it is imperative that we prioritise the conservation of these essential predators. This work also prompted several exciting follow-up considerations that could deepen our understanding of the intricate relationships between bats, birds, insects, and plant communities. For instance, exploring how these effects change in tropical forests, and understanding the impact of bottom-up control on the patterns found here. It was shown that the predation rate by ants strongly changes with increasing latitude, suggesting a potentially more pronounced impact in tropical regions (Zvereva et al., 2020), than what we have observed in this temperate study. Additionally, the effects of top-down and bottom-up forces on generalist and specialist herbivores differ, with bottom-up forces exerting stronger effects on specialists (Vidal & Murphy, 2018). Hence, future studies must consider the role of bottom-up forces to encompass a critical aspect of most ecological interactions.

## Notes

### Competing Interest Statement

The authors have declared no competing interest.

## References

Aceves-Aparicio, A., Narendra, A., McLean, D. J., Lowe, E. C., Christian, M., Wolff, J. O., Schneider, J. M., Herberstein, M. E. (2022). Fast acrobatic maneuvers enable arboreal spiders to hunt dangerous prey. Proceedings of the National Academy of Sciences, 119(40). 10.1073/pnas.2205942119

Aikens, K. R., Timms, L. L., & Buddle, C. M. (2013). Vertical heterogeneity in predation pressure in a temperate forest canopy. PeerJ, 1. 10.7717/peerj.138

Bael, S. A. V., Philpott, S. M., Greenberg, R., Bichier, P., Barber, N. A., Mooney, K. A., & Gruner, D. S. (2008). Birds as predators in tropical agroforestry systems. Ecology, 89(4), 928–934. 10.1890/06-1976.1

Bagchi, R., Gallery, R. E., Gripenberg, S., Gurr, S. J., Narayan, L., Addis, C. E., Freckleton, R. P., Lewis, O. T. (2014). Pathogens and insect herbivores drive rainforest plant diversity and composition. Nature, 506(7486), 85–88. 10.1038/nature12911

Balza, U., Lois, N. A., Polito, M. J., Pütz, K., Salom, A., & Raya Rey, A. (2020). The dynamic trophic niche of an island bird of prey. Ecology and evolution, 10(21), 12264–12276. 10.1002/ece3.6856

Barber, N. A., & Marquis, R. J. (2009). Spatial variation in top-down direct and indirect effects on white oak (Quercus alba L.). The American Midland Naturalist, 162(1), 169–179. 10.1674/0003-0031-162.1.169

Basset, Y. (2003). Arthropods of tropical forests: spatio-temporal dynamics and resource use in the canopy. Cambridge University Press.

Basset, Y., Cizek, L., Cuénoud, P., Didham, R. K., Novotny, V., Ødegaard, F., Roslin,T., Tishechkin, A. K., Schmidl, J., Winchester, N. N. (2015). Arthropod distribution in a tropical rainforest: tackling a four dimensional puzzle. Plos one, 10(12). 10.1371/journal.pone.0144110

Basset, Y., Hammond, P. M., Barrios, H., Holloway, J. D., & Miller, S. E. (2003). Vertical stratification of arthropod assemblages. Arthropods of tropical forests, 17–27. Cambridge University Press.

Bates, D., Kliegl, R., Vasishth, S., & Baayen, H. (2015). Parsimonious mixed models. arXiv preprint. 10.48550/arXiv.1506.04967

Beilke, E. A., & O’Keefe, J. M. (2023). Bats reduce insect density and defoliation in temperate forests: An exclusion experiment. Ecology, 104(2). 10.1002/ecy.3903

Belovsky, G., & Slade, J. (2000). Insect herbivory accelerates nutrient cycling and increases plant production. Proceedings of the National Academy of Sciences, 97(26), 14412–14417. 10.1073/pnas.250483797

Benitez-Lopez, A., Alkemade, R., Schipper, A., Ingram, D. J., Verweij, P., Eikelboom, J., & Huijbregts, M. (2017). The impact of hunting on tropical mammal and bird populations. Science, 356(6334), 180–183. 10.1126/science.aaj1891

Blaise, C., Mazzia, C., Bischoff, A., Millon, A., Ponel, P., & Blight, O. (2021). The key role of inter-row vegetation and ants on predation in Mediterranean organic vineyards. Agriculture, Ecosystems & Environment, 311. 10.1016/J.Agee.2021.107327

Blüthgen, N., & Feldhaar, H. (2010). Food and shelter: how resources influence ant ecology. Ant ecology, 115–136. 10.1093/acprof:oso/9780199544639.003.0007

Boege, K., & Marquis, R. J. (2006). Plant quality and predation risk mediated by plant ontogeny: consequences for herbivores and plants. Oikos, 115(3), 559–572. 10.1111/j.2006.0030-1299.15076.x

Böhm, S. M., Wells, K., & Kalko, E. K. (2011). Top-down control of herbivory by birds and bats in the canopy of temperate broad-leaved oaks (Quercus robur). Plos one, 6(4). 10.1371/journal.pone.0017857

Bolker, B., & Bolker, M. B. (2017). Package ‘bbmle’. Tools for General Maximum Likelihood Estimation, 641.

Bouarakia, O., Linden, V. M., Joubert, E., Weier, S. M., Grass, I., Tscharntke, T., Foord, S.H., Taylor, P. J. (2023). Bats and birds control tortricid pest moths in South African macadamia orchards. Agriculture, Ecosystems & Environment, 352. 10.1016/J.Agee.2023.108527

Brandt, M., & Mahsberg, D. (2002). Bugs with a backpack: the function of nymphal camouflage in the West African assassin bugs Paredocla andAcanthaspis spp. Animal Behaviour, 63(2), 277–284. 10.1006/anbe.2001.1910

Brooks, M. E., Kristensen, K., Van Benthem, K. J., Magnusson, A., Berg, C. W., Nielsen, A., Skaug, H. J., Machler, M., Bolker, B. M. (2017). glmmTMB balances speed and flexibility among packages for zero-inflated generalized linear mixed modeling. The R journal, 9(2), 378–400. 10.32614/Rj-2017-066

Brown, K. S. (1997). Diversity, disturbance, and sustainable use of Neotropical forests: insects as indicators for conservation monitoring. Journal of Insect conservation, 1(1), 25–42. 10.1023/A:1018422807610

Bulgarini, G., Castracani, C., Mori, A., Grasso, D. A., & Maistrello, L. (2021). Searching for new predators of the invasive Halyomorpha halys: the role of the black garden ant Lasius niger. Entomologia Experimentalis et Applicata, 169(9), 799–806. 10.1111/eea.13075

Cassano, C. R., Silva, R. M., Mariano-Neto, E., Schroth, G., & Faria, D. (2016). Bat and bird exclusion but not shade cover influence arthropod abundance and cocoa leaf consumption in agroforestry landscape in northeast Brazil. Agriculture, Ecosystems & Environment, 232, 247–253. 10.1016/j.agee.2016.08.013

Coley, P. D. (1991). Comparison of herbivory and plant defenses in temperate and tropical broad-leaved forest. Plant-animal interactions: evolutionary ecology in tropical and temperate regions, 25–49.

Coley, P. D., & Barone, J. (1996). Herbivory and plant defenses in tropical forests. Annual Review of Ecology and Systematics, 27(1), 305–335. 10.1146/annurev.ecolsys.27.1.305

Collins, J., & Jones, G. (2009). Differences in bat activity in relation to bat detector height: implications for bat surveys at proposed windfarm sites. Acta Chiropterologica, 11(2), 343–350. 10.3161/150811009X485576

Compton, S. G., Ellwood, M. D., Davis, A. J., & Welch, K. (2000). The flight heights of chalcid wasps (hymenoptera, Chalcidoidea) in a lowland Bornean rain forest: fig wasps are the high fliers 1. Biotropica, 32(3), 515–522. 10.1111/j.1744-7429.2000.tb00497.x

De Dijn, B. (2003). Vertical stratification of flying insects in a Surinam lowland rainforest. Arthropods of tropical forests, 110–122. Cambridge University Press.

De Vries, P. (1988). Stratification of fruit-feeding nymphalid butterflies in a Costa Rican rainforest. Journal of Research on the Lepidoptera, 26(1-4), 98–108.

Dekeukeleire, D., van Schrojenstein Lantman, I. M., Hertzog, L. R., Vandegehuchte, M. L., Strubbe, D., Vantieghem, P., Martel, A., Verheyen, K., Bonte, D., Lens, L. (2019). Avian top-down control affects invertebrate herbivory and sapling growth more strongly than overstorey species composition in temperate forest fragments. Forest ecology and management, 442, 1–9. 10.1016/j.foreco.2019.03.055

Del Hoyo, J., Elliot, A., & Sargatal, J. (1996). Handbook of the Birds of the World. Lynx Edicions.

Denmead, L. H., Darras, K., Clough, Y., Diaz, P., Grass, I., Hoffmann, M. P., Nurdiansyah, F., Fardiansah, R., Tscharntke, T. (2017). The role of ants, birds and bats for ecosystem functions and yield in oil palm plantations. Ecology, 98(7), 1945–1956. 10.1002/ecy.1882

DeVries, P. J., Murray, D., & Lande, R. (1997). Species diversity in vertical, horizontal, and temporal dimensions of a fruit-feeding butterfly community in an Ecuadorian rainforest. Biological Journal of the Linnean Society, 62(3), 343–364. 10.1111/j.1095-8312.1997.tb01630.x

Donald, P. F., Green, R., & Heath, M. (2001). Agricultural intensification and the collapse of Europe’s farmland bird populations. Proceedings of the Royal Society of London. Series B: Biological Sciences, 268(1462), 25–29. 10.1098/rspb.2000.1325

Eguchi, K. (1980). The feeding ecology of the nestling great tit, Parus major minor, in the temperate ever-green broadleaved forest II. With reference to breeding ecology. Researches on Population Ecology, 22(2), 284–300. 10.1007/BF02530852

Emlen, J. M. (1966). The role of time and energy in food preference. The American Naturalist, 100(916), 611–617. 10.1086/282455

Ferguson, K. I., & Stiling, P. (1996). Non-additive effects of multiple natural enemies on aphid populations. Oecologia, 108(2), 375–379. 10.1007/BF00334664

Ferreira, D. F., Jarrett, C., Wandji, A. C., Atagana, P. J., Rebelo, H., Maas, B., & Powell, L. L. (2023). Birds and bats enhance yields in Afrotropical cacao agroforests only under high tree-level shade cover. Agriculture, Ecosystems & Environment, 345. 10.1016/j.agee.2022.108325

Floren, A., Wetzel, W., & Staab, M. (2014). The contribution of canopy species to overall ant diversity (Hymenoptera: Formicidae) in temperate and tropical ecosystems. Myrmecological News, 19(March), 65–74.

Fukui, D., Agetsuma, N., & Hill, D. A. (2004). Acoustic identification of eight species of bat (Mammalia: Chiroptera) inhabiting forests of southern Hokkaido, Japan: potential for conservation monitoring. Zoological Science, 21(9), 947–955. 10.2108/zsj.21.947

Garcia, L. C., & Eubanks, M. D. (2019). Overcompensation for insect herbivory: a review and meta-analysis of the evidence. Ecology, 100(3). 10.1002/ecy.2585

Garrett, D. R., Pelletier, F., Garant, D., & Bélisle, M. (2022). Combined influence of food availability and agricultural intensification on a declining aerial insectivore. Ecological monographs, 92(3). https://doi.org/Artn E1518 10.1002/Ecm.1518

Gossner, M. M., Pašalić, E., Lange, M., Lange, P., Boch, S., Hessenmöller, D., Müller, J., Socher, S. A., Fischer, M., Schulze, E.-D. (2014). Differential responses of herbivores and herbivory to management in temperate European beech. Plos one, 9(8). 10.1371/journal.pone.0104876

Gras, P., Tscharntke, T., Maas, B., Tjoa, A., Hafsah, A., & Clough, Y. (2016). How ants, birds and bats affect crop yield along shade gradients in tropical cacao agroforestry. Journal of Applied Ecology, 53(3), 953–963. 10.1111/1365-2664.12625

Greenberg, R., Bichier, P., Angon, A. C., MacVean, C., Perez, R., & Cano, E. (2000). The impact of avian insectivory on arthropods and leaf damage in some Guatemalan coffee plantations. Ecology, 81(6), 1750–1755. 10.2307/177321

Haack, N., Borges, P. A., Grimm-Seyfarth, A., Schlegel, M., Wirth, C., Bernhard, D., Brunk, I., Henle, K., Pereira, H. M. (2022). Response of common and rare beetle species to tree species and vertical stratification in a floodplain forest. Insects, 13(2), 161. 10.3390/insects13020161

Hill, C. J., Gillison, A. N., & Jones, R. E. (1992). The spatial distribution of rain forest butterflies at three sites in North Queensland, Australia. Journal of Tropical Ecology, 8(1), 37–46. 10.1017/S0266467400006064

Hölldobler, B., & Wilson, E. O. (1990). The ants. Harvard University Press.

Holmes, R. T., Schultz, J. C., & Nothnagle, P. (1979). Bird predation on forest insects: an exclosure experiment. Science, 206(4417), 462–463. 10.1126/science.206.4417.462

Chapman, S. K., Hart, S. C., Cobb, N. S., Whitham, T. G., & Koch, G. W. (2003). Insect herbivory increases litter quality and decomposition: an extension of the acceleration hypothesis. Ecology, 84(11), 2867–2876. 10.1890/02-0046

Ichinose, K. (1990). The Ant Fauna of the Tomakomai Experiment Forest, Hokkaido University (Hymenoptera: Formicidae) with Notes on the Nuptial Season. Research Bulletins of College Experiment Forests, 47(1), 137–144.

Imai, H. T., Kihara, A., Kondoh, M., Kubota, M., Kuribayashi, S., Ogata, K., Onoyama, K., Taylor, R. W., Terayama, M.,Tsukii, Y., Yoshimura, M., Ugawa, Y. (2003). Ants of Japan. Gakken.

Intachat, J., & Holloway, J. (2000). Is there stratification in diversity or preferred flight height of geometroid moths in Malaysian lowland tropical forest? Biodiversity & Conservation, 9(10), 1417–1439. 10.1023/A:1008926814229

Janzen, D. H. (1970). Herbivores and the number of tree species in tropical forests. The American Naturalist, 104(940), 501–528. 10.1086/282687

Johnson, M., Kellermann, J., & Stercho, A. (2010). Pest reduction services by birds in shade and sun coffee in Jamaica. Animal conservation, 13(2), 140–147. 10.1111/j.1469-1795.2009.00310.x

Kalcounis, M., Hobson, K., Brigham, R., & Hecker, K. (1999). Bat activity in the boreal forest: importance of stand type and vertical strata. Journal of Mammalogy, 80(2), 673–682. 10.2307/1383311

Kalka, M., & Kalko, E. K. (2006). Gleaning bats as underestimated predators of herbivorous insects: diet of Micronycteris microtis (Phyllostomidae) in Panama. Journal of Tropical Ecology, 22(1), 1–10. 10.1017/S0266467405002920

Karp, D. S., & Daily, G. C. (2014). Cascading effects of insectivorous birds and bats in tropical coffee plantations. Ecology, 95(4), 1065–1074.

Kerbiriou, C., Bas, Y., Le Viol, I., Lorrilliere, R., Mougnot, J., & Julien, J. F. (2019). Potential of bat pass duration measures for studies of bat activity. Bioacoustics, 28(2), 177–192. 10.1080/09524622.2017.1423517

Kollross, J., Jancuchova-Laskova, J., Kleckova, I., Freiberga, I., Kodrik, D., & Sam, K. (2023). Nonlethal Effects of Predation: The Presence of Insectivorous Birds (Parus major) Affects The Behavior and Level of Stress in Locusts (Schistocerca gregaria). Journal of Insect Behavior, 36(1), 68–80. 10.1007/s10905-023-09820-z

Larrivée, M., & Buddle, C. M. (2009). Diversity of canopy and understorey spiders in north- temperate hardwood forests. Agricultural and Forest Entomology, 11(2), 225–237. 10.1111/j.1461-9563.2008.00421.x

Lenth, R. (2018). Emmeans: Estimated marginal means, aka least-squares means (Version R Package Version 1.2.4.).

Liefke, C., Dorow, W., Hölldobler, B., & Maschwitz, U. (1998). Nesting and food resources of syntopic species of the ant genus Polyrhachis (Hymenoptera, Formicidae) in West-Malaysia. Insectes sociaux, 45(4), 411–425. 10.1007/s000400050098

Lichtenberg, J. S., & Lichtenberg, D. A. (2002). Weak trophic interactions among birds, insects and white oak saplings (Quercus alba). The American Midland Naturalist, 148(2), 338–349. 10.1674/0003-0031(2002)148[0338:Wtiabi]2.0.Co;2

Losey, J. E., & Denno, R. F. (1998). Positive predator–predator interactions: enhanced predation rates and synergistic suppression of aphid populations. Ecology, 79(6), 2143–2152. 10.2307/176717

Lüdecke, D., Ben-Shachar, M. S., Patil, I., Waggoner, P., & Makowski, D. (2021). performance: An R package for assessment, comparison and testing of statistical models. Journal of Open Source Software, 6(60). 10.21105/joss.03139

Maas, B., Clough, Y., & Tscharntke, T. (2013). Bats and birds increase crop yield in tropical agroforestry landscapes. Ecology Letters, 16(12), 1480–1487. 10.1111/ele.12194

Maas, B., Heath, S., Grass, I., Cassano, C., Classen, A., Faria, D., Gras, P., Williams-Guillén, K., Johnson, M., Karp, D. S. (2019). Experimental field exclosure of birds and bats in agricultural systems—Methodological insights, potential improvements, and cost-benefit trade-offs. Basic and Applied Ecology, 35, 1–12. 10.1016/j.baae.2018.12.002

MacArthur, R. H., & Pianka, E. R. (1966). On optimal use of a patchy environment. The American Naturalist, 100(916), 603–609. 10.1086/282454

Maguire, D. Y., Nicole, T., Buddle, C. M., & Bennett, E. M. (2015). Effect of fragmentation on predation pressure of insect herbivores in a north temperate deciduous forest ecosystem. Ecological Entomology, 40(2), 182–186. 10.1111/een.12166

Marra, P. P., & Remsen Jr, J. (1997). Insights into the maintenance of high species diversity in the Neotropics: habitat selection and foraging behavior in understory birds of tropical and temperate forests. Ornithological Monographs, 445–483. 10.2307/40157547

Matsuo, T., Hiura, T., & Onoda, Y. (2022). Vertical and horizontal light heterogeneity along gradients of secondary succession in cool-and warm-temperate forests. Journal of Vegetation Science, 33(3). 10.1111/jvs.13135

Mestre, L., Piñol, J., Barrientos, J., Cama, A., & Espadaler, X. (2012). Effects of ant competition and bird predation on the spider assemblage of a citrus grove. Basic and Applied Ecology, 13(4), 355–362. 10.1016/j.baae.2012.04.002

Mols, C. M., & Visser, M. E. (2002). Great tits can reduce caterpillar damage in apple orchards. Journal of Applied Ecology, 39(6), 888–899. 10.1046/j.1365-2664.2002.00761.x

Mooney, K. A. (2007). Tritrophic effects of birds and ants on a canopy food web, tree growth, and phytochemistry. Ecology, 88(8), 2005–2014. 10.1890/06-1095.1

Mooney, K. A., Gruner, D. S., Barber, N. A., Van Bael, S. A., Philpott, S. M., & Greenberg, R. (2010). Interactions among predators and the cascading effects of vertebrate insectivores on arthropod communities and plants. Proceedings of the National Academy of Sciences, 107(16), 7335–7340. 10.1073/pnas.1001934107

Morrison, E. B., & Lindell, C. A. (2012). Birds and bats reduce insect biomass and leaf damage in tropical forest restoration sites. Ecological Applications, 22(5), 1526–1534. 10.1890/11-1118.1

Murakami, M. (2002). Foraging mode shifts of four insectivorous bird species under temporally varying resource distribution in a Japanese deciduous forest. Ornithological Science, 1(1), 63–69. 10.2326/osj.1.63

Nakamura, A., Kitching, R. L., Cao, M., Creedy, T. J., Fayle, T. M., Freiberg, M., Hewitt, C. N., Itioka, T., Koh, L. P, Ma, K. (2017). Forests and their canopies: achievements and horizons in canopy science. Trends in ecology & evolution, 32(6), 438–451. 10.1016/j.tree.2017.02.020

Nyffeler, M., Şekercioğlu, Ç. H., & Whelan, C. J. (2018). Insectivorous birds consume an estimated 400–500 million tons of prey annually. The Science of Nature, 105, 1–13. 10.1007/s00114-018-1571-z

Ocampo-Ariza, C., Vansynghel, J., Bertleff, D., Maas, B., Schumacher, N., Ulloque- Samatelo, C., Yovera, F. F., Evert, T., Ingolf, S. D., Tscharntke, T. (2023). Birds and bats enhance cacao yield despite suppressing arthropod mesopredation. Ecological Applications, 33(5). 10.1002/eap.2886

Offenberg, J. (2001). Balancing between mutualism and exploitation: the symbiotic interaction between Lasius ants and aphids. Behavioral Ecology and Sociobiology, 49(4), 304–310. 10.1007/s002650000303

Ohyama, L., King, J. R., & Jenkins, D. G. (2020). Are tiny subterranean ants top predators affecting aboveground ant communities? Ecology, 101(8). 10.1002/ecy.3084

Ozanne, C. M., Anhuf, D., Boulter, S. L., Keller, M., Kitching, R. L., Körner, C., Meinzer, F. C., Mitchell, A. W., Nakashizuka, T., Dias, P. L., Stork, N. E., Wright, S. J., Yoshimura, M., Dias, P. S. (2003). Biodiversity meets the atmosphere: a global view of forest canopies. Science, 301(5630), 183–186. 10.1126/science.1084507

Paine, R. T. (1966). Food web complexity and species diversity. The American Naturalist, 100(910), 65–75. 10.1086/282400

Paine, R. T. (1980). Food webs: linkage, interaction strength and community infrastructure. Journal of Animal Ecology, 49(3), 667–685. 10.2307/4220

Parker, G. G. (1995). Structure and microclimate of forest canopies. Forest canopies., 73–106. Academic press.

Pérez-Espona, S. (2021). Eciton Army ants—Umbrella species for conservation in neotropical forests. Diversity, 13(3), 136. 10.3390/d13030136

Perfecto, I., & Vandermeer, J. (1996). Microclimatic changes and the indirect loss of ant diversity in a tropical agroecosystem. Oecologia, 108(3), 577–582. 10.1007/BF00333736

Philpott, S. M., & Armbrecht, I. (2006). Biodiversity in tropical agroforests and the ecological role of ants and ant diversity in predatory function. Ecological Entomology, 31(4), 369–377. 10.1111/j.1365-2311.2006.00793.x

Philpott, S. M., Greenberg, R., Bichier, P., & Perfecto, I. (2004). Impacts of major predators on tropical agroforest arthropods: comparisons within and across taxa. Oecologia, 140(1), 140–149. 10.1007/s00442-004-1561-z

Philpott, S. M., Perfecto, I., & Vandermeer, J. (2008). Effects of predatory ants on lower trophic levels across a gradient of coffee management complexity. Journal of Animal Ecology, 77(3), 505–511. 10.1111/j.1365-2656.2008.01358.x

Piel, G., Tallamy, D. W., & Narango, D. L. (2021). Lepidoptera host records accurately predict tree use by foraging birds. Northeastern Naturalist, 28(4), 527–540. 10.1656/045.028.0410

Plank, M., Fiedler, K., & Reiter, G. (2012). Use of forest strata by bats in temperate forests. Journal of Zoology, 286(2), 154–162. 10.1111/j.1469-7998.2011.00859.x

Reynolds, B. C., & Crossley Jr, D. (1997). Spatial variation in herbivory by forest canopy arthropods along an elevation gradient. Environmental Entomology, 26(6), 1232–1239. 10.1093/Ee/26.6.1232

Richards, L. A., & Coley, P. D. (2007). Seasonal and habitat differences affect the impact of food and predation on herbivores: a comparison between gaps and understory of a tropical forest. Oikos, 116(1), 31–40. 10.1111/j.2006.0030-1299.15043.x

Rosumek, F. B., Silveira, F. A., de S. Neves, F., de U. Barbosa, N. P., Diniz, L., Oki, Y., Pezzini, F., Fernandes, G. W., Cornelissen, T. (2009). Ants on plants: a meta-analysis of the role of ants as plant biotic defenses. Oecologia, 160(3), 537–549. 10.1007/s00442-009-1309-x

Sam, K., Jorge, L. R., Koane, B., Amick, P. K., & Sivault, E. (2023). Vertebrates, but not ants, protect rainforest from herbivorous insects across elevations in Papua New Guinea. Journal of biogeography, 50(10), 1803–1816. 10.1111/jbi.14686

Sam, K., Koane, B., Bardos, D. C., Jeppy, S., & Novotny, V. (2019). Species richness of birds along a complete rain forest elevational gradient in the tropics: Habitat complexity and food resources matter. Journal of biogeography, 46(2), 279–290. 10.1111/jbi.13482

Sam, K., Koane, B., Sam, L., Mrazova, A., Segar, S., Volf, M., Moos, M., Simek, P., Sisol, M., Novotny, V. (2020). Insect herbivory and herbivores of Ficus species along a rain forest elevational gradient in Papua New Guinea. Biotropica, 52(2), 263–276. 10.1111/btp.12741

Sanders, D., & van Veen, F. F. (2011). Ecosystem engineering and predation: the multi- trophic impact of two ant species. Journal of Animal Ecology, 80(3), 569–576. 10.1111/j.1365-2656.2010.01796.x

Sayama, K., Ito, M., Tabuchi, K., Ueda, A., Ozaki, K., & Hironaga, T. (2012). Seasonal trends of forest moth assemblages in Central Hokkaido, Northern Japan. The Journal of the Lepidopterists’ Society, 66(1), 11–26. 10.18473/lepi.v66i1.a2

Schemske, D. W., Mittelbach, G. G., Cornell, H. V., Sobel, J. M., & Roy, K. (2009). Is there a latitudinal gradient in the importance of biotic interactions? Annual Review of Ecology, Evolution, and Systematics., 40, 245–269. 10.1146/annurev.ecolsys.39.110707.173430

Schifani, E., Castracani, C., Giannetti, D., Spotti, F. A., Reggiani, R., Leonardi, S., Mori, A., Grasso, D. A. (2020). New tools for conservation biological control: Testing ant-attracting artificial Nectaries to employ ants as plant defenders. Insects, 11(2), 129. 10.3390/Insects11020129

Schulze, C. H., Linsenmair, K. E., & Fiedler, K. (2001). Understorey versus canopy: patterns of vertical stratification and diversity among Lepidoptera in a Bornean rain forest. Plant Ecology, 153(1),133–152. 10.1023/A:1017589711553

Seifert, B. (2008). The ants of Central European tree canopies (Hymonoptera: Formicidae)-an underestimated population? Canopy arthropod research in Europe, In Floren, A., & Schmidl, J. (Eds.), Canopy arthropod research in Europe (pp. 157–173). Bioform Verlag.

Sih, A., Englund, G., & Wooster, D. (1998). Emergent impacts of multiple predators on prey. Trends in ecology & evolution, 13(9), 350–355. 10.1016/s0169-5347(98)01437-2

Singer, M. S., Clark, R. E., Lichter-Marck, I. H., Johnson, E. R., & Mooney, K. A. (2017). Predatory birds and ants partition caterpillar prey by body size and diet breadth. Journal of Animal Ecology, 86(6), 1363–1371. 10.1111/1365-2656.12727

Sivault, E., Amick, P. K., Armstrong, K. N., Novotny, V., & Sam, K. (2023). Species richness and assemblages of bats along a forest elevational transect in Papua New Guinea. Biotropica, 55(1), 81–94. 10.1111/btp.13161

Stephens, P. A., Mason, L. R., Green, R. E., Gregory, R. D., Sauer, J. R., Alison, J., Aunis, A., Brotons, L., Brutchart, S. H., M., Campedelli, T. (2016). Consistent response of bird populations to climate change on two continents. Science, 352(6281), 84–87. 10.1126/science.aac4858

Team, R. C. (2020). R: a language and environment for statistical computing. Version 4.0. 2. Vienna, Austria. In.

Thomine, E., Jeavons, E., Rusch, A., Bearez, P., & Desneux, N. (2020). Effect of crop diversity on predation activity and population dynamics of the mirid predator Nesidiocoris tenuis. Journal of Pest Science, 93(4), 1255–1265. 10.1007/s10340-020-01222-w

Thurman, J. H., Northfield, T. D., & Snyder, W. E. (2019). Weaver ants provide ecosystem services to tropical tree crops. Frontiers in ecology and evolution, 7, 120. 10.3389/Fevo.2019.00120

Tobing, M. C., & Kuswardani, R. A. (2018). The potential of Myopopone castanea (Hymenoptera: Formicidae) as a predator for Oryctes rhinoceros Linn. larvae (Coleoptera: Scarabaeidae). Journal of Physics: Conference Series, 1116(5). 10.1088/1742-6596/1116/5/052074

Tuma, J., Eggleton, P., & Fayle, T. M. (2020). Ant-termite interactions: an important but under-explored ecological linkage. Biological reviews, 95(3), 555–572. 10.1111/brv.12577

Ulyshen, M. D. (2011). Arthropod vertical stratification in temperate deciduous forests: implications for conservation-oriented management. Forest ecology and management, 261(9), 1479–1489. 10.1016/j.foreco.2011.01.033

Van Bael, S. A., Brawn, J. D., & Robinson, S. K. (2003). Birds defend trees from herbivores in a Neotropical forest canopy. Proceedings of the National Academy of Sciences, 100(14), 8304–8307. 10.1073/pnas.1431621100

Verboven, N., Tinbergen, J. M., & Verhulst, S. (2001). Food, reproductive success and multiple breeding in the great tit Parus major. Ardea, 89(2), 387–406.

Vidal, M. C., & Murphy, S. M. (2018). Bottom-up vs. top-down effects on terrestrial insect herbivores: A meta-analysis. Ecology Letters, 21(1), 138–150. 10.1111/ele.12874

Wang, X.-F., Liu, J.-F., Gao, W.-Q., Deng, Y.-P., Ni, Y.-Y., Xiao, Y.-H., Kang, F.-F., Wang, Q., Lei, J.-P., Jiang, Z.-P. (2016). Defense pattern of Chinese cork oak across latitudinal gradients: influences of ontogeny, herbivory, climate and soil nutrients. Scientific reports, 6(1), 27269. 10.1038/srep27269

Wesołowski, T., Rowiński, P., & Neubauer, G. (2019). Food of Nuthatch Sitta europaea young in a primeval forest: effects of varying food supply and age of nestlings. Acta Ornithologica, 54(1), 85–104. 10.3161/00016454AO2019.54.1.008

Williams-Guillén, K., Perfecto, I., & Vandermeer, J. (2008). Bats limit insects in a neotropical agroforestry system. Science, 320(5872), 70–70.

Wilson, E. O. (2000). Sociobiology: The new synthesis. Harvard University Press.

Wu, L., Kato, T., Sato, H., Hirano, T., & Yazaki, T. (2019). Sensitivity analysis of the typhoon disturbance effect on forest dynamics and carbon balance in the future in a cool-temperate forest in northern Japan by using SEIB-DGVM. Forest ecology and management, 451. 10.1016/J.Foreco.2019.117529

Zachos, F. E. (2020). DE Wilson and RA Mittermeier (chief editors): Handbook of the Mammals of the World. Vol. 9. Bats. Springer.

Zeus, V. M., Puechmaille, S. J., & Kerth, G. (2017). Conspecific and heterospecific social groups affect each other’s resource use: a study on roost sharing among bat colonies. Animal Behaviour, 123, 329–338. 10.1016/j.anbehav.2016.11.015

Zvereva, E. L., Zverev, V., & Kozlov, M. V. (2020). Predation and parasitism on herbivorous insects change in opposite directions in a latitudinal gradient crossing a boreal forest zone. Journal of Animal Ecology, 89(12), 2946–2957. 10.1111/1365-2656.13350

