## Supporting information for "Insectivorous birds and bats outperform ants in the top-down regulation of arthropods across strata of a Japanese temperate forest"

**APPENDIX S1**

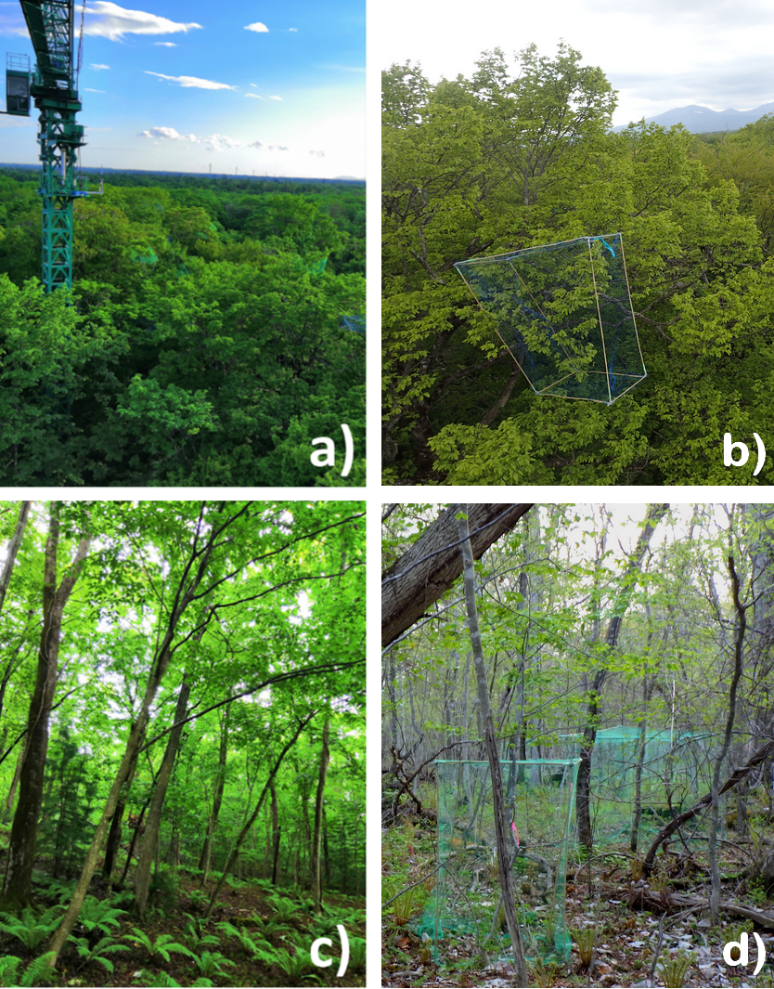

Figure S1.1:  Photos depicting (a) the characteristic forest canopy (with several detectable vertebrate exclosures) (c) the understory in the study area (b) the vertebrate exclosure treatment attached within the canopy and (d) the vertebrate exclosure treatments situated in the understory. Photographed by Jan Kollross.

Table S1.1:  Plant species used in the UBB (understory species) and the CUVA experiments (both understory and canopy species).

| Family | Plant species | Understory | Canopy |
| --- | --- | --- | --- |
| Magnoliaceae | *Magnolia kobus* | X | X |
| Oleaceae* | *Syringa reticulata* | X |  |
| Betulaceae | *Carpinus cordata* | X | X |
| Sapindaceae | *Acer mono* | X | X |
| Rosaceae | *Prunus ssiori* | X | X |
| Sapindaceae | *Acer palmatum* | X | X |
| Oleaceae | *Fraxinus lanuginosa* | X | X |
| Betulaceae** | *Betula maximowicziana* | X |  |
| Betulaceae** | *Ostrya japonica* |  | X |

*Note that *Syringa reticulata* was absent from the canopy, resulting in a difference in the number of tree species between the canopy and understory in the CUVA experiment.
**Note that *Betula maximowicziana* never grew taller than 3 m in the experimental forest, and *Ostrya japonica* rarely occurred as reachable saplings. Therefore, these two species were paired for the purpose of the CUVA experiment.

Table S1.2: Study design in CUVA experiment. Overview of the treatments within the experiment, their location, number of species and individuals used in each of them and experiment duration.

| Strata | Treatment | Species | Individuals per species and treatment | Duration (days) |
| --- | --- | --- | --- | --- |
| Understory | VER–exclusion of vertebrates | 8 | 5 | 65 ± 4 |
| Understory | ALL- exclusion of ants and vertebrates | 8 | 5 | 65 ± 4 |
| Understory | ANT- exclusion of ants | 8 | 5 | 65 ± 4 |
| Understory | CN2- control | 8 | 5 | 65 ± 4 |
| Canopy | VER– exclusion of vertebrates | 7 | 3 | 65 ± 4 |
| Canopy | ALL- exclusion of ants and vertebrates | 7 | 3 | 65 ± 4 |
| Canopy | ANT- exclusion of ants | 7 | 3 | 65 ± 4 |
| Canopy | CN2-control | 7 | 3 | 65 ± 4 |

Table S1.3: Study design in UBB experiment. Overview of the treatments within the experiment, their location, exclosure manipulation, number of species and individuals used in each of them and experiment duration.

| Strata | Treatment | Exclosure closed | Species | Individuals per species and treatment | Duration (days) |
| --- | --- | --- | --- | --- | --- |
| Understory | BIR – exclusion of birds | Night | 8 | 5 | 30 ± 2 |
| Understory | BAT- exclusion of bats | Day | 8 | 5 | 30 ± 2 |
| Understory | CN1- control | - | 8 | 5 | 30 ± 2 |

Table S1.4: Starting and ending dates of both experiments (CUVA and UBB experiments) in the years 2018 and 2019.

| Experiment | Year | Start | End |
| --- | --- | --- | --- |
| UBB | 2018 | 20-23.5. | 19-22.6. |
|  | 2019 | 12-15.5. | 11-14.6. |
| CUVA | 2018 | 12-22.5. | 22-31.7. |
|  | 2019 | 17.5.-8.6. | 20-29.7. |

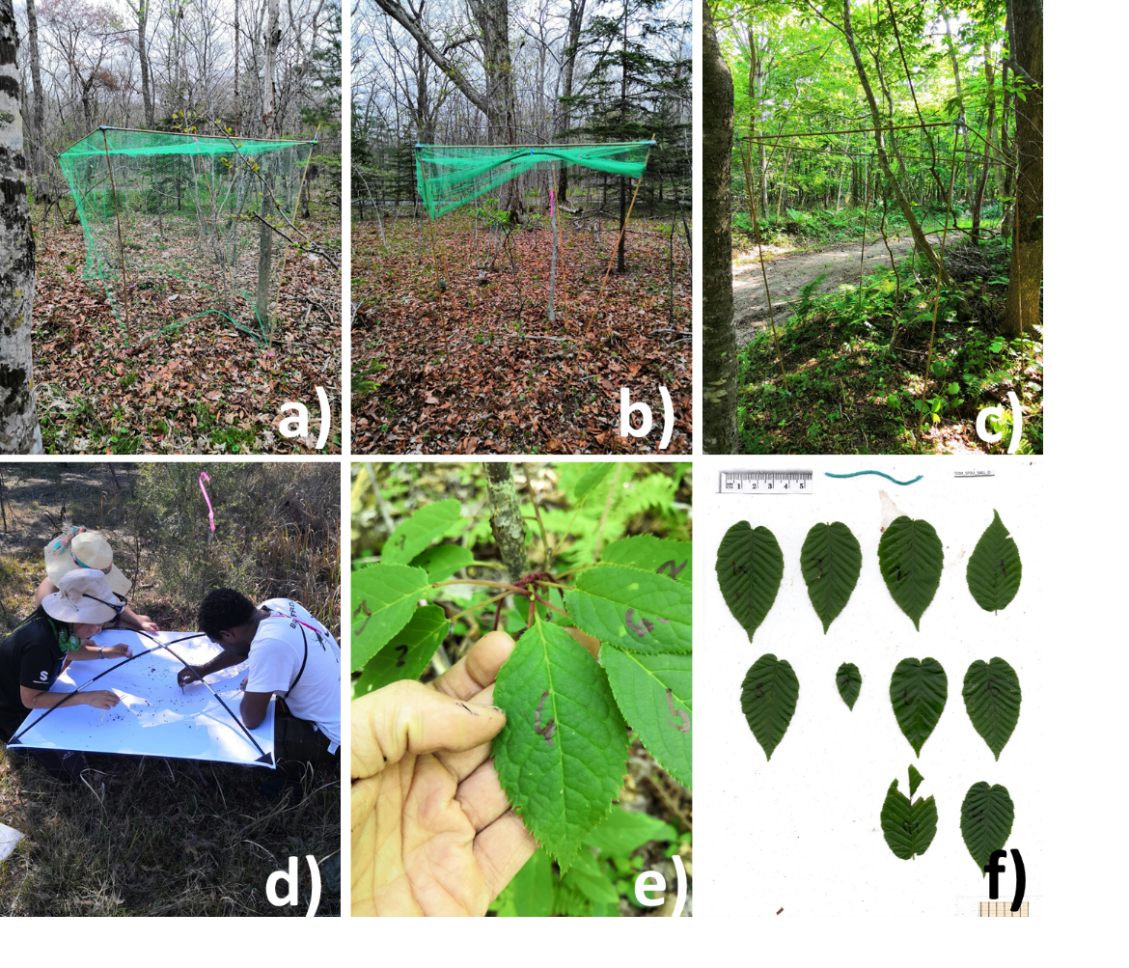

Figure S1.2:  Photos depicting (a) the vertebrate exclosure treatment in the forest understory, (b) the bird and bat exclosure treatment, with the mesh pulled up allowing the entrance of birds during the day (or bats during the night), (c) the FCN2 construction treatment, (d) the collection of the arthropods from the beating sheet at the end of the experiment, (e) the red string marked twig with numbered leaves ready for the collection at the end of the experiment and (f) the scanning of the leaves at the end of the experiment. Photographed by Jan Kollross and Elise Sivault.

**APPENDIX S2**

Table S2.1: Comparisons of multi-predictor models analysing factors (i.e., treatment and strata) affecting the density of all arthropods and herbivory damage, in the  CUVA experiment. Results of the analysis of deviance based on the delta AICc (Corrected Akaike Information Criterion). The most parsimonious models are indicated in bold, and the selected models are indicated in grey. Note that the second-best model has been selected for arthropod densities due to multicollinearity issues.

|  | Arthropod densities | | Herbivory damage | |
| --- | --- | --- | --- | --- |
|  | dAICc | df | dAICc | df |
| Null | 90.3 | 5 | 91.2 | 6 |
| Treatment | 61.5 | 8 | 37.8 | 9 |
| Strata | 68.9 | 6 | 61.6 | 7 |
| Strata + Plant species:Strata | 35.4 | 8 | 60.6 | 9 |
| Treatment + Strata | 38.1 | 9 | 3.1 | 10 |
| Treatment + Strata + Plant species:Strata | 0.5 | 11 | **0.0** | **12** |
| Treatment * Strata | 37.6 | 12 | 6.1 | 13 |
| Treatment * Strata + Plant species:Strata | **0.0** | **14** | 3.1 | 15 |

Table S2.2: Comparisons of multi-predictor models analysing factors  (i.e, treatment and strata) affecting the density of leaf chewers, predators, sapsuckers and NR (i.e., arthropod with no consumptive effect on other arthropods or plants) in the CUVA experiment, based on the delta AICc (Corrected Akaike Information Criterion). The most parsimonious models are indicated in bold and the selected models to plot the graphics are indicated in grey. Note that the selected model for chewer density had to contain the variable treatment for visualisation purposes (Figure 4).

|  | Chewer densities | | Predator densities | | Sapsucker densities | | NR densities | |
| --- | --- | --- | --- | --- | --- | --- | --- | --- |
|  | dAICc | df | dAICc | df | dAICc | df | dAICc | df |
| Null | **0.0** | **5** | 21.5 | 5 | 23.6 | 5 | 5.7 | 5 |
| Treatment | 5.5 | 8 | 13.2 | 8 | 7.2 | 8 | 0.1 | 8 |
| Strata | 1..9 | 6 | 8.5 | 6 | 18.5 | 6 | 5.7 | 6 |
| Strata + Plant species: Strata | 4.0 | 8 | 9.5 | 8 | 17.9 | 8 | 8.4 | 8 |
| Treatment + Strata | 7.4 | 9 | **0.0** | **9** | 1.3 | 9 | **0.0** | **9** |
| Treatment + Strata + Plant species: Strata | 9.6 | 11 | 0.9 | 11 | **0.0** | **11** | 2.6 | 11 |
| Treatment* Strata | 9.2 | 12 | 5.8 | 12 | 1.3 | 12 | 4.2 | 12 |
| Treatment * Strata + Plant species: Strata | 11.5 | 14 | 6.7 | 14 | 0.1 | 14 | 6.8 | 14 |

Table S2.3: Estimated marginal means (=emmeans) of the CUVA experiment models including arthropod density and herbivory damage as response variables. Note that the arthropod density data were log-transformed prior to the analyses.

| Arthropod density | | | |
| --- | --- | --- | --- |
| Treatment-Strata | emmean | SE | df |
| ALL-Canopy | -6.19 | 0.173 | 13.4 |
| ANT-Canopy | -6.65 | 0.172 | 13.1 |
| CN2-Canopy | -6.62 | 0.172 | 13.1 |
| VER-Canopy | -6.02 | 0.172 | 13.1 |
| ALL-Understory | -5.55 | 0.249 | 11.2 |
| ANT-Understory | -6.02 | 0.249 | 11.2 |
| CN2-Understory | -5.98 | 0.249 | 11.2 |
| VER-Understory | -5.38 | 0.249 | 11.2 |
| Herbivory | | | |
| ALL-Canopy | 0.08 | 0.008 | 1327 |
| ANT-Canopy | 0.05 | 0.006 | 1327 |
| CN2-Canopy | 0.06 | 0.006 | 1327 |
| VER-Canopy | 0.08 | 0.009 | 1327 |
| ALL-Understory | 0.11 | 0.014 | 1327 |
| ANT-Understory | 0.08 | 0.010 | 1327 |
| CN2-Understory | 0.08 | 0.010 | 1327 |
| VER-Understory | 0.12 | 0.014 | 1327 |

Table S2.4: Estimated marginal means (=emmeans) of the CUVA and UBB experiment models including predator density as a response variable. Note that the data were log-transformed prior to the analyses.

| Predator density | | | |
| --- | --- | --- | --- |
| Treatment | emmean | SE | df |
| ALL | -8.45 | 0.429 | 31.4 |
| ANT | -9.63 | 0.426 | 30.4 |
| VER | -8.09 | 0.426 | 30.4 |
| CN2 | -8.60 | 0.426 | 30.4 |
| BIR | -6.95 | 0.405 | 28.2 |
| BAT | -7.24 | 0.405 | 28.2 |
| CN1 | -7.41 | 0.405 | 28.2 |

Table S2.5: Estimated marginal means (=emmeans) of the CUVA and UBB experiment models including chewer density as a response variable. Note that the data were log-transformed prior to the analyses.

| Chewer density | | | |
| --- | --- | --- | --- |
| Treatment | emmean | SE | df |
| ALL | -10.02 | 0.319 | 37.3 |
| ANT | -9.80 | 0.316 | 36.0 |
| VER | -9.77 | 0.316 | 36.0 |
| CN2 | -9.94 | 0.316 | 36.0 |
| BIR | -9.84 | 0.672 | 25.6 |
| BAT | -8.74 | 0.672 | 25.6 |
| CN1 | -8.89 | 0.672 | 25.6 |

Table S2.6: Estimated marginal means (=emmeans) of the CUVA and UBB experiment models including sapsucker density as a response variable. Note that the data were log-transformed prior to the analyses.

| Sapsucker density | | | |
| --- | --- | --- | --- |
| Treatment | emmean | SE | df |
| ALL | -9.73 | 0.307 | 74.9 |
| ANT | -10.90 | 0.303 | 71.5 |
| VER | -10.15 | 0.303 | 71.5 |
| CN2 | -11.53 | 0.303 | 71.5 |
| BIR | -8.37 | 0.796 | 20.1 |
| BAT | -10.02 | 0.796 | 20.1 |
| CN1 | -10.50 | 0.796 | 20.1 |

Table S2.7: Estimated marginal means (=emmeans) of the CUVA and UBB experiment models including NR density (i.e., arthropod with no consumptive effect on other arthropods or plants) as a response variable. Note that the data were log-transformed prior to the analyses.

| NR density | | | |
| --- | --- | --- | --- |
| Treatment | emmean | SE | df |
| ALL | -8.42 | 0.311 | 58.1 |
| ANT | -9.30 | 0.308 | 55.9 |
| VER | -8.44 | 0.308 | 55.9 |
| CN2 | -9.40 | 0.308 | 55.9 |
| BIR | -9.11 | 0.519 | 36.9 |
| BAT | -9.58 | 0.519 | 36.9 |
| CN1 | -10.93 | 0.519 | 36.9 |

Table S2.8: Results of the analysis of deviance examining the effect of the explanatory variable treatment on densities of all arthropods and herbivory damage, in the UBB experiment, based on the delta AICc (Corrected Akaike Information Criterion). The most parsimonious models are indicated in bold and the selected models to plot the graphics are indicated in grey.

|  | Arthropod densities | | Herbivory damage | |
| --- | --- | --- | --- | --- |
|  | dAICc | df | dAICc | df |
| Null | 11 | 4 | 4.9 | 5 |
| **Treatment** | **0** | **6** | **0.0** | **7** |

Table S2.9: Results of the analysis of deviance examining the effect of the explanatory variable treatment on densities of chewers, predators, sapsuckers and NR (i.e., arthropod with no consumptive effect on other arthropods or plants), in the UBB experiment, based on the delta AICc (Corrected Akaike Information Criterion). The most parsimonious models are indicated in bold and the selected models to plot the graphics are indicated in grey. Note that the selected model for chewer and predator densities had to contain the variable treatment for visualisation purposes (Figure 4).

|  | Chewer densities | | Predator densities | | Sapsucker densities | | NR densities | |
| --- | --- | --- | --- | --- | --- | --- | --- | --- |
|  | dAICc | df | dAICc | df | dAICc | df | dAICc | df |
| Null | **0.0** | **4** | **0.0** | **4** | 4.6 | 4 | 5 | 4 |
| Treatment | 1.4 | 6 | 3.1 | 6 | **0.0** | **6** | **0.0** | **6** |

 Table S2.10: Estimated marginal means (=emmeans) of the UBB experiment models including arthropod density and herbivory damage as response variables. Note that the arthropod density data were log-transformed prior to the analyses.

| Arthropod density | | | |
| --- | --- | --- | --- |
| Treatment | emmean | SE | df |
| BAT | -5.33 | 0.265 | 12.7 |
| BIR | -5.18 | 0.265 | 12.7 |
| CN1 | -5.82 | 0.265 | 12.7 |
| Herbivory | | | |
| BAT | 0.08276 | 0.009058 | 683 |
| BIR | 0.087143 | 0.009375 | 683 |
| CN1 | 0.067925 | 0.007715 | 683 |

**APPENDIX S3**

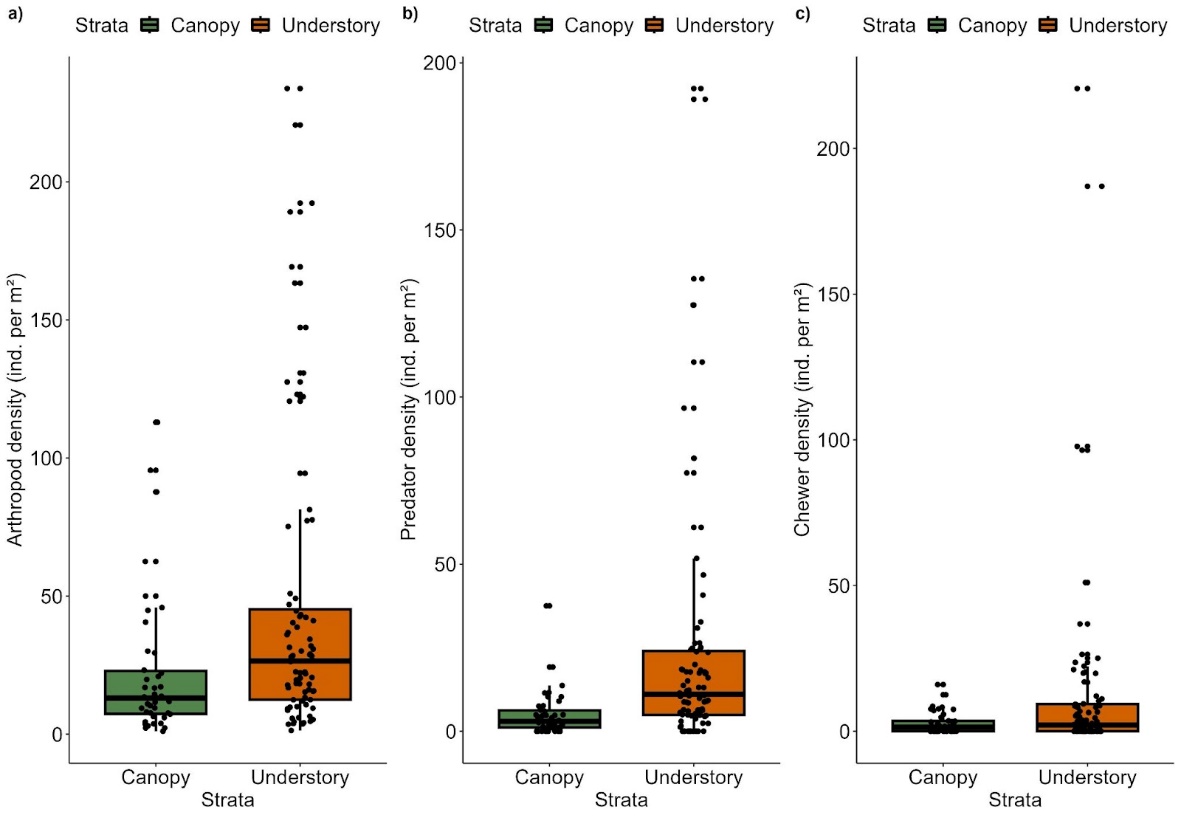

Figure S3.1: Total arthropod densities (individuals per square meter of foliage) (a), predator (b) and chewer (c) densities between canopy and understory on control treatments (CN2) in the CUVA experiment. The differences between the two strata are significant for the total arthropod and predator densities only (W=2245, P=0.002 and W=2539, P<0.001 respectively).

Table S3.1: Number of ants collected at the end of the CUVA experiment in each treatment (ANT, ALL, VER, and CON) at the canopy and understory levels. Note that zero ants were expected for ANT and ALL treatments if the removal of ants was 100% successful.

| Treatment | Canopy | Understory |
| --- | --- | --- |
| ANT | 1 | 13 |
| ALL | 2 | 36 |
| VER | 2 | 140 |
| CON | 2 | 100 |

Table S3.2: Percentages of increases or decreases of arthropod density partitioned into feeding guilds (NR: arthropod with no consumptive effect on other arthropods or plants) between treatments and experiments (canopy and understory combined). The significance is marked as follows: * P ≤ 0.05, ** P ≤ 0.01, *** P ≤ 0.001.

|  | Chewers | | Predators | | Sapsuckers | | NR | |
| --- | --- | --- | --- | --- | --- | --- | --- | --- |
|  | UBB | CUVA | UBB | CUVA | UBB | CUVA | UBB | CUVA |
| VER |  | 18 % |  | 66 % |  | 297 % *** |  | 161 %* |
| ALL |  | -7.6 % |  | 16 % |  | 504 % *** |  | 166 %* |
| ANT |  | 15 % |  | -64 %* |  | 87 % |  | 10 % |
| BIR | -61 % |  | 58 % |  | 741 %** |  | 517 %** |  |
| BAT | 16 % |  | 18 % |  | 61 % |  | 285 %* |  |

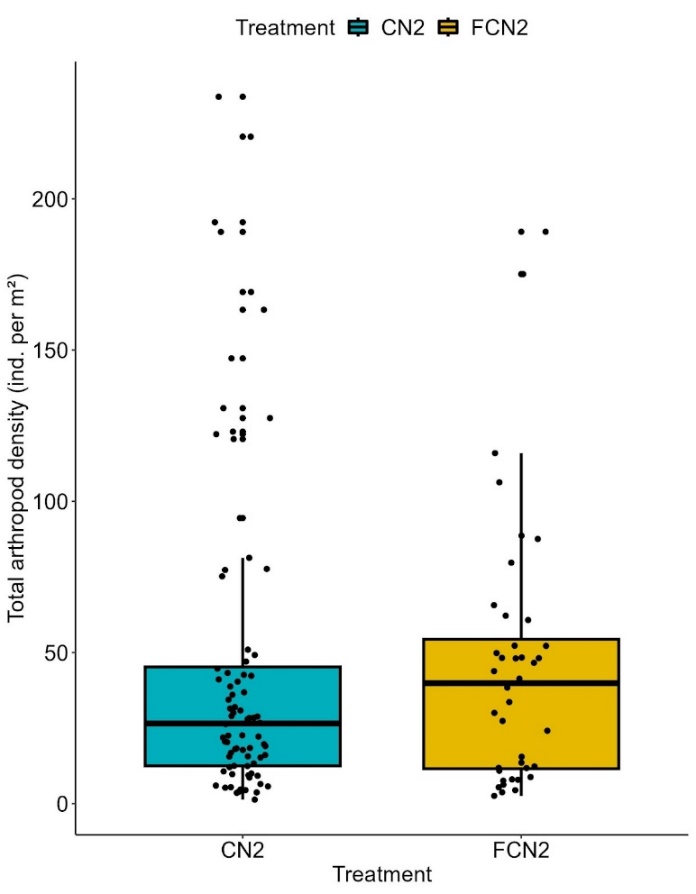

Figure S3.2: Total arthropod densities (individuals per square meter of foliage) between control (CN2) and frame-control (FCN2) in the understory (CUVA experiment). The difference in densities between the two controls is not significant (P>0.05).

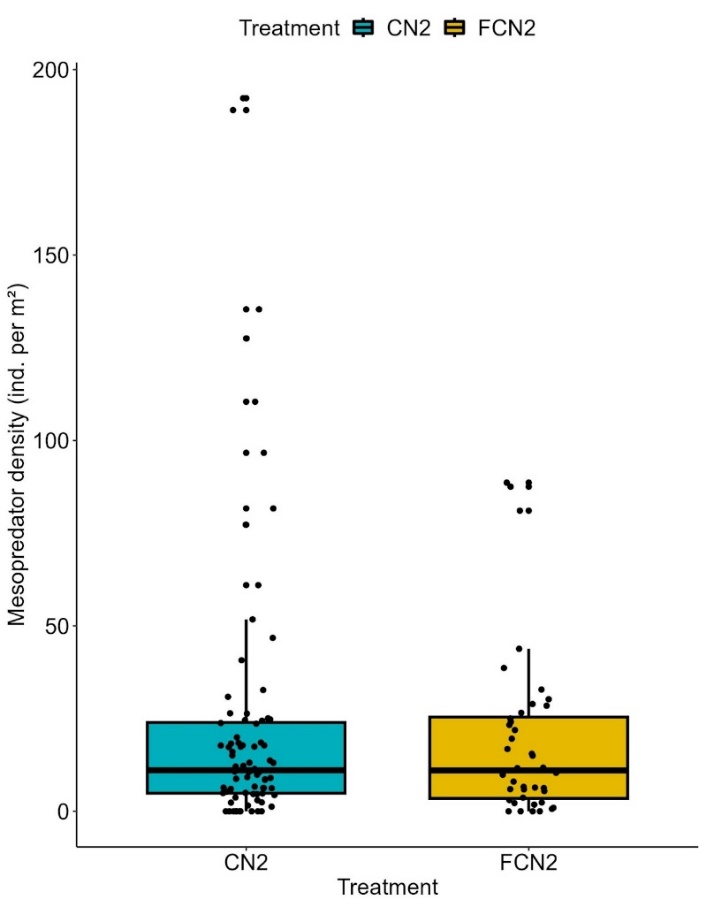

Figure S3.3: Mesopredator densities (individuals per square meter of foliage) between control (CN2) and frame-control (FCN2) in the understory (CUVA experiment). The difference in densities between the two controls is not significant (P>0.05).

Table S3.4: Results of the bird, bat, and ant surveys. For each species identified, abundance (ab.) or activity was estimated at each stratum. The body mass of the vertebrates (average or range) was found in the literature and used to calculate the biomass.

| **BIRDS** | | | |
| --- | --- | --- | --- |
| Species | Canopy ab. | Understory ab. | Weight (avg.) (g) |
| *Aegithalos caudatus* | 30 | 0 | 8.3 |
| *Anthus hodgsoni* | 8 | 1 | 21.6 |
| *Certhia familiaris* | 8 | 9 | 8.9 |
| *Cuculus optatus* | 22 | 6 | 106 |
| *Cyanoptila cyanomelana* | 0 | 9 | 25 |
| *Ficedula narcissina* | 89 | 63 | 11.5 |
| *Hierococcyx hyperythrus* | 2 | 0 | 120.1 |
| *Muscicapa dauurica* | 2 | 0 | 11.9 |
| *Paridae spp.* | 0 | 21 | 14.6 |
| *Parus minor* | 96 | 55 | 17 |
| *Periparus ater* | 93 | 20 | 9.6 |
| *Phylloscopus borealoides* | 17 | 0 | 10.7 |
| *Phylloscopus coronatus* | 203 | 24 | 9.2 |
| *Poecile palustris* | 54 | 10 | 11.9 |
| *Sitta europaea* | 25 | 3 | 22.5 |
| *Sittiparus varius* | 39 | 1 | 16.5 |
| *Turdidae sp.* | 0 | 1 | 70 |
| *Turdus cardis* | 5 | 22 | 65 |
| *Turdus chrysolaus* | 7 | 18 | 77 |
| *Urosphena squameiceps* | 0 | 63 | 9 |
| *Yungipicus kizuki* | 63 | 16 | 22 |
| *Zosterops japonicus* | 60 | 2 | 10.7 |
| **BATS** | | | |
| Species | Canopy ab. | Understory ab. | Weight (range) (g) |
| *Murina spp.* | 3 | 5 | 9-10.0 |
| *Myotis ikonnikovi* | 1 | 0 | 10 |
| *Myotis macrodactylus* | 0 | 6 | 6.0-8.0 |
| *Plecotus sacrimontis* | 1 | 0 | 7.4-9.2 |
| *Vespertilio sinensis* | 13 | 50 | 14-30 |
| **ANTS** | | | |
| Species | Canopy ab. | Understory ab. |  |
| *Aphaenogaster japonica* | 0 | 2 |  |
| *Camponotus obscuripes* | 0 | 9 |  |
| *Lasius hayashi* | 31 | 102 |  |
| *Lasius spathepus* | 24 | 48 |  |
| *Myrmica ruginodis* | 4 | 357 |  |
| *Nylanderia flavipes* | 2 | 281 |  |
| *Pheidole fervida* | 1 | 47 |  |
